## Supplementary Information for "Why is cancer not more common? A changing microenvironment may help to explain why, and suggests strategies for anti-cancer therapy"

#### Supplementary Figures

*Figure S1. Schematic view of the model as described in Methods. The whole model is shown in (**a**) and the chromosome instability model is shown in (**b**).*

*Figure S2. Examples of the adaptive cancer fitness landscapes used in this study.*

These examples illustrate how the fitness landscapes look like when different parameters are used. **a**-**d**, four different selection intensities are used in this study: $\sigma^2 = 10$ (**a**), $\sigma^2 = 40$ (**b**), $\sigma^2 = 70$ (**c**) and $\sigma^2 = 100$ (**d**). **e**-**f**, to illustrate how selections are correlated along traits in the fitness landscape two selection correlations are used (see equations (9) and Methods for more details): $\sigma^2 = 10$, $\rho_S = 0.9$ (**e**) and $\sigma^2 = 10$, $\rho_S = 0.5$ (**f**).

*Figure S3. Cancer evolutionary trajectories with a static TME optimum and different initial conditions.*

These results demonstrate that under a static TME optimum the initial fitness of cancer cell determines whether subsequent new driver mutations could be observed. If the initial fitness is high ($w_0 = 1$) there is no observation of any new driver mutation (**a**). If a lower fitness is assumed ($w_0 < 1$), there is a maximum of three driver mutations observed (**b**). However, if we assume the mutational variance $m^2 = 1 \times 10^{-5}$ and the selection intensity $\sigma^2 = 1$, the cancer with an initial fitness $w_0 = 0.9$ can adapt to the static TME optimum (**c**) by many "mini driver" mutations (**d**).

*Figure S4. Mean cancer cell population size under various selection intensities and phenotypic optimum change rates.*

Cancer populations evolve under four different rates of phenotypic optimum change ($v_1 = 0.05$, $v_1 = 5 \times 10^{-3}$, $v_1 = 5 \times 10^{-4}$ and $v_1 = 5 \times 10^{-5}$) with $\sigma^2 = 10$ (a), $\sigma^2 = 40$ (b),

$\sigma^2 = 70$ (c) and $\sigma^2 = 100$ (d). Error bars are s.e.m. and each point represents 100 independent simulations. The dash line represents population size at $N = 1 \times 10^6$.

*Figure S5. Mean cancer cell fitness under various selection intensities and TME conditions.*

These results indicate that selection intensity and phenotypic optimum change speed can significantly affect the evolutionary trajectories of a cancer. Cancer populations evolve under four different rates of phenotypic optimum change ($v_1 = 0.05$, $v_1 = 5 \times 10^{-3}$, $v_1 = 0.05$, $v_1 = 5 \times 10^{-4}$ and $v_1 = 5 \times 10^{-5}$) with $\sigma^2 = 10$ (**a**), $\sigma^2 = 40$ (**b**), $\sigma^2 = 70$ (**c**) and $\sigma^2 = 100$ (**d**). Error bars are s.e.m. and each point represents 100 independent simulations. The line represents a simple linear regression fit. Due to immediate population extinction data are not shown for $v_1 = 0.5$. The dash line represents mean fitness 0.5. When mean population fitness reaches this value it is destined to be extinct.

*Figure S6. Mean selection coefficients of adaptive mutations as per figure S5.*

These results indicate the selection intensity and phenotypic optimum changing speed play a significant role in cancer adaptive evolution.

*Figure S7. Effect of chromosome instability on cancer adaptation.*

The two plots show that too much chromosome instability is deleterious for cancer while the right amount can facilitate cancer adaptation when the TME selective pressure is high (e.g., high rate of phenotypic optimum change). The relation between chromosome instability rate and selection coefficient of selected driver mutations is shown in **a**. The relation between chromosome instability rate and mean population fitness is shown in **b**.

*Figure S8. Effect of mutation rate on cancer adaptation.*

These results show that when the phenotypic optimum changes increasing mutation rate can be beneficial to cancer as this can provide more beneficial mutations for selection to act on. Indeed, as the TME slows down this benefit becomes rather limited. The effect of two different mutation rates ($\mu = 4\times10^{-6}$ and $\mu = 4\times10^{-4}$) on adaptation is shown for selection coefficient of selected driver mutations in **a** and mean population fitness in **b**.

*Figure S9. Cancer adaptive evolution with different rates of selection correlation between traits.*

In a slowly changing TME the phenotypic effects of fixed mutations faithfully capture the selection correlation, which is defined by the fitness function (see equation (2) and Figure S2 for different shapes of the fitness landscape due different selection correlation values, defined by $\rho_S$). We have two traits here. The first trait, $\alpha_1$, is the trait that has a changing optimum, while the second trait, $\alpha_2$, has a contant phenotypic optium. Populations evolve under four different selection correlations and phenotypic optimum change rates. **a**, **e**, **i** and **m**, $\rho_S = 0.25$. **b**, **f**, **j** and **n**, $\rho_S = 0.5$. **c**, **g**, **k** and **o**, $\rho_S = 0.75$. **d**, **h**, **l** and **p**, $\rho_S = 0.9$.

*Figure S10. Modulating TME selection through cancer cell-TME interaction.*

TME selection is modulated by modeling selection intensity as a robustness parameter with a cost to the cancer cell. These simulations are performed in a directionally changing TME with $v_1 = 5\times10^{-3}$, $v_1 = 5\times10^{-4}$ and $v_1 = 5\times10^{-5}$, respectively. The relation between mean population fitness and robustness is fitted with a simple linear model. The population is evolved under low ($N = 1\times10^{6}$, blue line) and high ($N = 1\times10^{3}$, red line) genetic drift, respectively.

*Figure S11. Mean fitness and selection coefficient of cancer adaptation in a randomly changing TME.*

These results show that the increased variance in random changes of the TME acts against adaptive cancer evolution, which leads to decreased mean population fitness and increased selection coefficients of fixed mutations. The mean fitness (**a**) and selection coefficient (**b**) are plotted against different standard deviations (SD) of the random phenotypic optimum change. Error bars are the standard error of the mean (s.e.m.), and each point represents 100 independent simulations.

Figure S12. *Mean fitness and selection coefficient of cancer adaptation in a directionally changing TME with a random component.*

Here it is similar to Figure S11 that increased variance in the directionally changing TME acts against adaptive cancer evolution. The mean fitness (**a**) and selection coefficient (**b**) are plotted against different standard deviations (SD) of the random change. Different colours represent different speed of the directional change. Error bars are the standard error of the mean (s.e.m.), and each point represents 100 independent simulations.

Figure S13. *Mean fitness and selection coefficient of cancer adaptation in a cyclically changing TME.*

These results suggest that when the period of the cyclically changing TME is fixed the increased amplitude can decrease mean population fitness and increase the selection coefficients of fixed mutations, which act against adaptive cancer evolution. The mean fitness (**a**) and selection coefficient (**b**) are plotted against different amplitudes of the phenotypic optimum change. The period is set at $P = 360$. Error bars are the standard error of the mean (s.e.m.), and each point represents 100 independent simulations.

*Figure S14. Sub-clonal competition initiated by mini drivers in a directionally changing TME.*

The ancestral clone is initiated with 10 cells, while the two sub-clones are both started with 2 cells. Each parameter combination (**a**-**c**) is performed 100 times and sampled every 10 generations. The two neutral mutant sub-clones are shown in (**a**) for easy comparison. The two sub-clones initiated by two driver mutations with equal competition are shown in (**b**), while the two sub-clones with stronger competition are shown in (**c**). Three different summary statistics are shown, namely, mean population size (N), mean population fitness (w), and variance of the mean fitness (w-var).

*Figure S15. Sub-clonal competition initiated by intermediate drivers in a directionally changing TME as per Figure S14.*

*Figure S16. Sub-clonal competition initiated by classic major drivers in a directionally changing TME as per Figure S14.*

*Figure S17. Sub-clonal competition initiated by mini drivers in a randomly changing TME as per Figure S14.*

*Figure S18. Sub-clonal competition initiated by intermediate drivers in a randomly changing TME as per Figure S14.*

*Figure S19. Sub-clonal competition initiated by classic major drivers in a randomly changing TME as per Figure S14.*

*Figure S20. Sub-clonal competition initiated by mini drivers in a cyclically changing TME as per Figure S14.*

*Figure S21. Sub-clonal competition initiated by intermediate drivers in a cyclically changing TME as per Figure S14.*

*Figure S22. Sub-clonal competition initiated by classic major drivers in a cyclically changing TME as per Figure S14.*

*Figure S23. Properties of genotypic landscapes generated from Fisher's phenotypic landscapes with a changing TME.*

The simulations are performed in a changing TME with three different rates of the phenotypic optimum ($v_1 = 5\times10^{-3}$, $v_1 = 5\times10^{-4}$ and $v_1 = 5\times10^{-5}$) using three ancestral fitness, 0.1 (**a**), 0.5 (**b**) and 0.9 (**c**), respectively. Each parameter combination is simulated 100 times. In each simulation we calculated the epistasis among selected driver mutations, the fraction of sign epistasis and the roughness to slope ratio with different rates of phenotypic optimum change following Methods. We fitted the results with linear models to show the general rations between these parameters and phenotypic optimum change rates.

*Figure S24. Relation between roughness to slop ratio, epistasis and the fraction of sign epistasis as per Figure S23.*

We use the same simulations as in Figure S23 to demonstrate the simple relation between the fraction of sign epistasis and roughness to slope ratio. Both can be used to characterize the ruggedness of the underlying genotypic landscapes generated from Fisher's phenotypic landscape.

*Figure S25. Cancer phylogenies under randomly and cyclically changing TME selection dynamics*

Example phylogenetic trees are shown for simulated cancers under two different TME selective dynamics. **a-d**, four phylogenetic trees are shown for a randomly changing TME with four different standard deviations: $\delta = 0.5$ (**a**), $\delta = 1$ (**b**), $\delta = 1.5$ (**c**) and $\delta = 2$ (**d**). **e-h**, four phylogenetic trees are shown for a cycling TME with four different amplitudes: $A = 0.5$ (**e**), $A = 2$ (**f**), $A = 4$ (**g**) and $A = 6$ (**h**). All cancers were longitudinally sampled for every 100 generations for a fixed period of time of 10000 generations. The maximum population size is set at $N = 10^5$. The scale bar represents the number of cell divisions.

*Figure S26. Cancer adaptation with different spatial constraints and changing TMEs.*

These results show that increasing the 3D space size of the TME can facilitate adaptation especially when the phenotypic optimum changes fast. These simulations are performed with three different optimum changing dynamics: directional (a, d), random (b, e) and cyclic (c, f). The maximum population size is set at $N = 1 \times 10^5$ for all simulations. The 3D space size for spatial constraints is set at $300^3$ (green), $200^3$ (red) and $100^3$ (blue), respectively

*Figure S27. Quantifying phylogeny asymmetry under different spatial constraints and changing TMEs.*

The measure of phylogeny asymmetry with the number of cherries and the normalized Sackin's index can clearly reflect the tumour's underlying evolutionary dynamics. When the phenotypic optimum changes fast, the increased number of cherries in larger TME space indicate large population size during each sampling interval (**a-c**), which could make selection more effective. The increased normalized Sackin's index further support this (**d-f**). These simulations are performed as per Figure S26.

*Figure S28. Illustration of the fitness landscape shifts due to different anti-cancer treatment strategies.*

These figures illustrate how the optimum of a fitness landscape moves quantitatively and how the reduced selection intensity leads to a "flatter" fitness landscape in cancer treatments. **a**, fitness landscape changes with different (constant) TME optimum due to different treatments. **b**, the optimum of the fitness landscape changes from $z_0^{opt} = 0$ to $z_1^{opt} = 5$. **c**, fitness landscape optimum changes from $z_0^{opt} = 0$ to $z_1^{opt} = 8$. **d**, fitness landscape optimum changes from $z_0^{opt} = 0$ to $z_1^{opt} = 8$ and selection intensity from $\sigma^2 = 10$ to $\sigma^2 = 40$. Clearly, the shorter distance that an optimum travels gives a less "steep" valley for the cancer cells to cross (evolve resistance). Similarly, the reduced selection intensity (a "flatter" fitness landscape) also gives the cancer cells a less "steep" valley to cross from the original fitness landscape to the new one (compare **c** and **d**).

*Figure S29. Cancer evolution under different anti-cancer treatment strategies.*

Example 3D snapshots are taken sequentially after sudden change of the TME optimum due to treatments (**a-o**). Four different treatments maintaining four different levels of TME optimum are shown: $z_1^{opt} = 5$ (**a-e**), $z_1^{opt} = 6$ (**f-j**), $z_1^{opt} = 7$ (**k-o**) and $z_1^{opt} = 8$ (no 3D snapshots are taken due to immediate population extinction after treatment), respectively. For understanding clonality of cancer cells after treatments, **e**, **j** and **o** show the clonal expansion of cancer cells after treatments for $z_1^{opt} = 5$, $z_1^{opt} = 6$ and $z_1^{opt} = 7$, respectively. In order to track the precise fitness status of the population the sample is taken for every generation. The population fitness plotted against generation time is summarized for each treatment (**p**), and the dashed line indicates when the treatment starts (after generation 100). We first assume that the cancer has evolved to a constant TME optimum $z_0^{opt} = 0$ and allow it 100 generations to accumulate genetic variation and reach maximum tumour size ($N = 1\times10^7$). We then assess four different TME optima to represent different treatment strategies ($z_1^{opt} = 5,6,7,8$, see equation (17), Supplementary Movies S22-S25) that treat the cancer for about 33 months or more (e.g., 1000 generations), reducing mean population fitness below $w = 0.1$. We illustrate how the optimum of the fitness landscape changes from $z_0^{opt}$ to $z_1^{opt}$ under each treatment (Supplementary Figure S28). All treatments reduce the fitness of all cancers below $w = 0.1$ (**a-o**) and at $z_1^{opt} = 8$ the cancer is successfully cured and the population is extinct (**p**), but due to mutation and smaller phenotypic change (e.g., because of smaller dose or effectiveness of the delivery) the cancers survive at $z_1^{opt} = 5,6,7$ and quickly relapse in less than two months (**p**).

*Figure S30. Cancer adaptation with different resistance mechanisms under treatment strategies maintaining a constant TME optimum.*

Different resistance mechanisms lead to different 3D spatio-temporal patterns of sub-clonal evolution. Example 3D snapshots are taken sequentially after the sudden change of TME optimum ($z_1^{opt} = 8, \sigma^2 = 10$) due to treatments (**a-d**), however, the population has three different resistance mechanisms to avoid population extinction. **a**, the cancer cells have more loci contributing to adaptation ($L = 50$). We find two adaptive mutations at generation 113 and generation 1033, which are born in generation 101 and 228 with $s = 38.281$ and $s = 0.5976$, respectively. **b**, the cancer cells have higher mutation rates ($\mu = 4\times10^{-4}$). In this case we find one adaptive mutation at generation 977 born at generation 101 with a very large selection coefficient, $s = 57.3361$, indicating very strong selection leading to the fixation of this mutation with very large fitness effect. **c**, the cancer cells evolve in a fitness landscape with lower selection intensity ( $\sigma^2 = 40$ ). The population fitness is summarized in **d** for each treatment strategy.

*Figure S31. Cancer adaptation under treatment strategies that continuously change the TME optimum.*

Treatment strategies continuously modify the TME optimum leading to different 3D patterns of spatio-temporal patterns of sub-clonal evolution and evolutionary trajectories. Example 3D snapshots are taken sequentially after treatments (**a-d**). Three different treatments maintaining three different rates of phenotypic optimum change: $v_1 = 0.5$ (**a**), $v_1 = 0.05$ (**b**) and $v_1 = 0.005$ (**c**), respectively. The population fitness plotted against generation time is summarized in **d** for each treatment strategy. Note that the sample is taken for every generation, and the dashed line indicates when the treatment starts (after generation 100).

#### Supplementary Movies

Movie S1. A simulation movie showing 3D cancer adaptation under a static TME with $v_1 = 0$.

Movie S2. A simulation movie showing 3D cancer adaptation under a phenotypic optimum change rate $v_1 = 0.5$.

Movie S3. A simulation movie showing 3D cancer adaptation under a phenotypic optimum change rate $v_1 = 0.05$.

Movie S4. A simulation movie showing 3D cancer adaptation under a phenotypic optimum change rate $v_1 = 5 \times 10^{-3}$.

Movie S5. A simulation movie showing 3D cancer adaptation under a phenotypic optimum change rate $v_1 = 5 \times 10^{-4}$.

Movie S6. A simulation movie showing 3D cancer adaptation under a phenotypic optimum change rate $v_1 = 5 \times 10^{-5}$.

Movie S7. A simulation movie showing 3D cancer adaptation under a phenotypic optimum change rate $v_1 = 5 \times 10^{-5}$. In this simulation we have initial fitness $w = 0.1$, initial population size $N = 10^4$.

Movie S8. A simulation movie showing 3D cancer adaptation under a phenotypic optimum change rate $v_1 = 0.05$, initial fitness $w = 0.1$, initial population size $N = 10^7$.

Movie S9. A simulation movie showing 3D cancer adaptation under a phenotypic optimum change rate $v_1 = 5 \times 10^{-3}$, initial fitness $w = 0.1$, initial population size $N = 10^7$.

Movie S10. A simulation movie showing 3D cancer adaptation under a phenotypic optimum change rate $v_1 = 5\times10^{-4}$, initial fitness $w = 0.1$, initial population size $N = 10^7$.

Movie S11. A simulation movie showing 3D cancer adaptation under a phenotypic optimum change rate $v_1 = 5\times10^{-5}$, initial fitness $w = 0.1$, initial population size $N = 10^7$.

Movie S12. A simulation movie showing 3D cancer adaptation under a phenotypic optimum change rate $v_1 = 0.05$, initial fitness $w = 0.5$, initial population size $N = 10^7$.

Movie S13. A simulation movie showing 3D cancer adaptation under a phenotypic optimum change rate $v_1 = 5\times10^{-3}$, initial fitness $w = 0.5$, initial population size $N = 10^7$.

Movie S14. A simulation movie showing 3D cancer adaptation under a phenotypic optimum change rate $v_1 = 5\times10^{-4}$, initial fitness $w = 0.5$, initial population size $N = 10^7$.

Movie S15. A simulation movie showing 3D cancer adaptation under a phenotypic optimum change rate $v_1 = 5\times10^{-5}$, initial fitness $w = 0.5$, initial population size $N = 10^7$.

Movie S16. A simulation movie showing 3D cancer adaptation under a cyclically changing TME. The amplitude is set at $A = 4$ with period $P = 360$.

Movie S17. A simulation movie showing 3D cancer adaptation under a cyclically changing phenotypic optimum. The amplitude is set at $A = 6$ with period $P = 360$.

Movie S18. A simulation movie showing 3D sub-clonal competition under a randomly changing TME. The two mutant sub-clones are initiated by two neutral mutations (red and yellow), respectively. This simulation shows that a neutral mutant sub-clone can dominate the tumour and become fixed, which suggests the changing TME dynamics plays an important role in determining the evolutionary trajectories of sub-clones. The standard deviation of the randomly changing TME is set at $\delta = 1.5$.

Movie S19. A simulation movie showing 3D sub-clonal competition under a cyclically changing TME. The two mutant sub-clones are initiated by two mini drivers (red with 1% selective advantage and yellow with 5% selective advantage), respectively. This simulation shows that one of the sub-clones is lost due to clonal competition while the other sub-clone becomes dominant and fixed. The TME amplitude is set at $A = 2$ with period $P = 360$.

Movie S20. A simulation movie showing 3D sub-clonal competition under a randomly changing TME. The two mutant sub-clones are initiated by two intermediate drivers both with 10% selective advantage (red and yellow), respectively. This simulation shows that the two sub-clones can coexist for an extremely long period of time. The standard deviation of the changing TME is set at $\delta = 1.5$.

Movie S21. A simulation movie showing 3D sub-clonal competition under a randomly changing TME. The two mutant sub-clones are initiated by two major classic drivers both with 20% selective advantage (red and yellow), respectively. This simulation shows that although the two sub-clones have large selective advantage, due to strong competition and a random TME both sub-clones went extinct eventually. The standard deviation of the changing TME is set at $\delta = 1$.

Movie S22. A simulation movie showing 3D cancer adaptation under a treatment strategy maintaining a constant TME optimum $z_1^{opt} = 5$.

Movie S23. A simulation movie showing 3D cancer adaptation under a treatment strategy maintaining a constant TME optimum $z_1^{opt} = 6$.

Movie S24. A simulation movie showing 3D cancer adaptation under a treatment strategy maintaining a constant TME optimum $z_1^{opt} = 7$.

Movie S25. A simulation movie showing 3D cancer adaptation under a treatment strategy maintaining a constant TME optimum $z_1^{opt} = 8$. Note that in this simulation the cancer population went extinct.

Movie S26. A simulation movie showing 3D cancer adaptation with a resistance mechanism under a treatment strategy maintaining a constant TME optimum at $z_1^{opt} = 8$. The resistance mechanism here is the number of loci contributing to adaptation is increased to $L = 50$ from $L = 5$. Comparing to Movie S25, the population extinction is avoided.

Movie S27. A simulation movie showing 3D cancer adaptation with a resistance mechanism under a treatment strategy maintaining a constant TME optimum at $z_1^{opt} = 8$. The resistance mechanism here is increased mutation rate from $\mu = 4 \times 10^{-5}$ to $\mu = 4 \times 10^{-4}$. Comparing to Movie S25, the population extinction is also avoided.

Movie S28. A simulation movie showing 3D cancer adaptation with a resistance mechanism under a treatment strategy maintaining a constant TME optimum at $z_1^{opt} = 8$. The resistance mechanism here is that the TME selection intensity is reduced from $\sigma^2 = 10$ to $\sigma^2 = 40$ (the width is increased see Supplementary Figure S2a and 2b). Comparing to Movie S25, the population extinction is also avoided.

### Supplementary Notes

In the Supplementary Notes 1-3, we investigate how other model parameters could have affected cancer adaptation. First, we find that different chromosome instability rates, mutation rates and number of cancer traits can affect cancer adaptation to various degrees (Supplementary Figures S7-S8, Supplementary Note 1). Particularly, we show that only the right amount of chromosome instability, the "just-right" model, can facilitate cancer adaption when the TME selection is harsh and too much chromosome instability is indeed deleterious. Second, we demonstrate that the pleiotropic driver mutations and selection correlation can lead to systematic maladaptation among cancer traits and therefore hamper cancer adaptation (Supplementary Figure S9, Supplementary Note 2), which could be exploited for treatment purposes. Third, our further model extension shows that TME selection could be modulated through cancer cell-TME interaction with an adjustable fitness cost (Supplementary Figure S10, Supplementary Note 3).

#### Supplementary Note 1

##### Potential effects of chromosome instability, mutation and cancer hallmark traits on cancer adaptation in a changing TME

We seek to understand how chromosome instability rate, mutation rate and the number of cancer traits affect cancer adaptation. First, for simplicity we only performed the same simulations shown in the results section, where the cancer cell population starts from an optimum phenotype with seven different chromosome instability rates. The results are consistent with the notion that too much chromosome instability is deleterious and the right amount can help tumour's adaptation when the environmental selective pressure is strong[1]. This pattern can be clearly shown when the chromosome instability rate increases there is a clear decrease of mean tumour fitness when the TME selective pressure is strong (v=0.05, Supplementary Figure S7). Second, when the per locus mutation rate is increased from $4\times10^{-6}$ to $4\times10^{-4}$, there is indeed an increase of mean cancer cell population fitness, accompanied by a reduction in mean selective advantage of driver mutations (Supplementary Figure S8). Third, as we increase the number of cancer cell traits (from 1 dimension to 8 dimensions, see Methods), we find that there is a cost of complexity associated with adaptive cancer evolution. In particular, the mean fitness of the population decreases and the mean selective advantage increases during adaptation when the cancer cell has an increased number of traits and changing TME (Figure 2d-e). This observation could be understood in terms of the hallmark traits of cancer cells[2]. With increase number of hallmark traits, the rate of cancer population adaptation could be reduced. However, if some of the eight hallmark traits act in the same dimension through modularity the cost could be reduced but can not be eliminated[3]. As a result, the cancer with lower phenotypic complexity may persist for longer period of time and it is more likely to establish a clinically significant cancer.

#### Supplementary Note 2

#### Effects of pleiotropic driver mutations, selection correlation and maladaptation among cancer hallmark traits

In cancer evolution, most master driver mutations have pleiotropic effects, a single genotype affecting multiple pathways and several cancer traits. Although we assumed that each mutation affects all traits (universal pleiotropy), its fitness effect on each trait may be different. In other words, the driver mutations may have different fitness effects on different traits and the selection is therefore correlated (equation (2)), which can be illustrated by the shapes of the fitness landscapes with different levels of correlation (see Supplementary Figure S2). Similarly, a mutation (genotype) may have different phenotypic effects on each trait. So, the mutational effects on traits can be independent or correlated (similar to equation (2)). Indeed, we find that when the phenotypic optimum changes slowly, the fitness effects of driver mutations along traits indeed correlate (Supplementary Figure S9). When adapting to the first trait with a changing optimum selection correlation causes systematic maladaptation of cancer population in the second trait (this trait has a constant optimum at the origin), which can be clearly seen in a linear fasion (Supplementary Figure S9). Moreover, we find that when the phenotypic optimum changes fast the phenotypic effects of driver mutations along traits have similar levels of correlation as correlations of mutational effects (data not shown). Although in clinical settings the effects of selectional and mutational correlations on cancer progression are unclear, an example may be multiple mutations with pleiotropic effect on multiple cancer phenotypic traits in Wnt signalling pathway genes (APC, TCF7L2, SOX9) in colorectal cancer, or multiple pleiotropic mutations in Pi3K genes (e.g. KRAS and PIK3CA) in several cancers.

#### Supplementary Note 3

#### Modulating TME selection through cancer cell-TME interaction

As shown above we did not consider cancer cell-TME interactions. However, there might be a feedback/interplay between cancer cells and their TME affecting a cancer's evolutionary trajectory. We therefore model the TME selection intensity as the genetic robustness of cancer cells with a fitness cost [4], where lowering TME selection intensity and/or increasing the robustness of the cancer cell's response will come at a fitness cost to the cancer cells. Briefly, we introduce a cost parameter in the fitness function in the simple form of

$$w(\mathbf{z},t) = \frac{1}{C(r^2)} \exp\left[ -\left(\mathbf{z} - \mathbf{z}^{opt}(t)\right)^T R^{-1} \left(\mathbf{z} - \mathbf{z}^{opt}(t)\right) \right].$$

Here $C\left(r^2\right)$ is a cost function and $r^2$ is the robustness ( $r^2 \geq 1$). The matrix $\mathbf{R}$ is a real $n \times n$ positive definite and symmetrical matrix and can be defined similarly as $\mathbf{S}$ in equation (2). To understand the effect of robustness, we perform simulations under different rates of phenotypic optimum changes. Moreover, we consider whether the cancer cells evolving under low or high genetic drift would make a difference (Drift is generally linked to population size). In general, high robustness is deleterious to cancer cells as this can lead to high cost to the cancer cells (Supplementary Figure S10), so there is a trade-off here. However, it is more deleterious to cancer cells evolving in a slowly changing TME than in a fast changing TME. Interestingly, when the phenotypic optimum change speed is low, increasing robustness can be more deleterious under low genetic drift (Supplementary Figure S10).

#### Supplementary Note 4

##### Cancer adaptation in randomly and cyclically changing TMEs

First, when the phenotypic optimum changes randomly, the resulting increase in its variance at different time points always acts against cancer adaptation (equations (6)-(8)), leading to reduced mean population fitness and a requirement for driver mutations with higher mean fitness effect (Supplementary Figure S11). Second, when we add a random component into a directionally changing TME (equations (9)-(11)), the increased variance caused by the random component also acts against cancer adaptation, producing similar results to a purely randomly changing TME (Supplementary Figure S12). Third, when the phenotypic optimum changes cyclically (equations (12)-(14)), with increased amplitude, the cycling TME optimum also acts against cancer adaptation (Supplementary Figure S13). However, interestingly, although the mean population fitness decreases and mean selective advantage of selected driver mutations increase when the amplitude increases, there are more selected driver mutations recorded than under any other phenotypic optimum change dynamics (full data not shown, amplitude $A=4$, see Supplementary Figure S13 and Supplementary Movies S16-S17). Moreover, there are also more complex spatio-temporal patterns of sub-clonal fitness and mixing, in which birth and death of large and small sub-clones with diverse fitness values occur frequently through time and space (Supplementary Movies S16-S17). This indicates that a cycling TME at intermediate level may be particularly capable of promoting cancer adaptation and allow cancer cells to reach their phonotypic optimum by periodically fixing more driver mutations than TMEs that change directionally and/or randomly.

#### Supplementary Note 5

#### Sub-clonal competition initiated by driver mutations with various selective advantages in a changing TME

We have shown that due to clonal interference there are multiple beneficial mutant sub-clones competing in the population in a changing TME. Moreover, neutral mutant sub-clones may become fixed and dominate the population. To understand these sub-clonal evolutionary dynamics in a changing TME, we have extended our model to address sub-clonal evolution explicitly, based on one ancestral clone and two mutant sub-clones competing with each other. We model four types sub-clonal competitions in three types of changing TME, namely, directional (Supplementary Figures S14-S16), random (Supplementary Figures S17-S19) and cycling TME (Supplementary Figures S20-S22). We first simulate the evolution of neutral mutants (sub-clones initiated by mutations with no selective advantage, $s=0$) and use neutrality as a baseline, and then we compare this to the sub-clones initiated by three different types of driver mutations we show above: mini ($s=1\%$), intermediate ($s=10\%$) and classic ($s=20\%$)[5]. The two mutant sub-clones can have equal initial fitness advantage (initiated by driver mutations of the same selective advantage) or one of the sub-clones can be a competitor initiated by a driver mutation with a relatively higher fitness advantage. We set these sub-clonal competitors with initial driver mutations of selective advantage at: mini (sub-clone 1, $s_1=1\%$ vs sub-clone 2, $s_2=5\%$), intermediate (sub-clone 1, $s_1=5\%$ vs sub-clone 2, $s_2=10\%$) and classic ( sub-clone 1, $s_1=20\%$ vs sub-clone 2, $s_2=30\%$ ). In all cases, we show clonal interference to various degrees leading to the loss of one of the advantageous sub-clones. But their evolutionary trajectories are still determined by how their phenotypic optimum changes, which follow similar evolutionary dynamics in a changing TME as shown above. However, we reveal the details of sub-clonal competition not seen above. First, we find that neutral mutants generally become extinct quickly. Nevertheless, due to the stochastic nature of a random TME, neutral mutant sub-clones could become fixed and dominate the tumour (Supplementary Movie S18). Second, mutant sub-clones initiated by mini drivers can also become fixed (Supplementary Movie S19), although not necessarily dominating early cancer. Third, sub-clones with intermediate drivers can easily dominate early cancer and become fixed. Interestingly, a randomly changing TME can maintain the coexistence of two advantageous sub-clones for a long period of time (Supplementary Movie S20). Finally, sub-clones with classic major drivers can both dominate and become fixed in early cancer. Strikingly, due to strong competition, occasionally both advantageous sub-clones can become extinct (Supplementary Movie S21).

Understanding how caner evolution and ecology couples to affect cancer dynamics is of paramount importance to develop next-generation cancer therapies[6-8]. Here, our modelling framework attempts to offer an integrated view. We show that the sub-clonal evolutionary trajectories initiated by cell-autonomous driver mutations in early tumours are generally consistent with the size of their fitness effect and relative population size[9]. However, the competitions between multiple advantageous sub-clones could weaken the efficiency of selection and alter these trajectories. Moreover, the changing TME dynamics can also play a critical role in this process. For instance, a randomly changing TME can lead to complete disappearance of two advantageous sub-clones with a selective advantage as high as 20%. Moreover, it can maintain the coexistence of two intermediately advantageous sub-clones for a long period of time. A random TME can also lead a neutral mutant sub-clone to dominate the tumour and become fixed. Nevertheless, the non-cell autonomous role of these constantly changing TME selective dynamics in determining the sub-clonal evolutionary trajectories are generally not considered in previous studies[9-12], which may lead to considerable biases in inferring the underlying cancer dynamics.

#### Supplementary Note 6

#### Properties of genotypic fitness landscapes of selected driver mutations generated by Fisher's phenotypic fitness landscape with a changing phenotypic optimum

To show that Fisher's phenotypic landscape can generate various genotypic landscapes, we use our modelling framework with two types of changing TME (directional and cyclic) to generate genotypic landscapes of selected driver mutations (we use four driver mutations, see Methods). We first calculate all pairwise epistasis coefficients among selected driver mutations and find that all epistasis coefficients are generally negative indicating the combined effect of driver mutations is smaller than their independent effects. Negative epistasis is important in explaining why driver mutations may acquire early, which is consistent with previous studies in microbial evolution experiments that the rate of adaptation tends to slow down over time [13]. We then calculate the fraction of sign epistasis (simple and/or complex sign epistasis wherever possible) and roughness to slope ratio (see Methods). Interestingly, all changing TMEs lead to extensive sign epistasis and roughness of the underlying genotypic landscapes of driver mutations. For instance, we show that increasing the rate of directionally changing TME can increase the ruggedness of the underlying genotypic landscapes, which is manifested as the increased fraction of sign epistasis and roughness to slope ratio (Supplementary Figures 23-24). The rate of phenotypic optimum change can also have a negative impact on the epistasis coefficient of selected driver mutations (Supplementary Figure 24). These results suggest the rather smooth Fisher's phenotypic landscape with a changing TME can generate Wright's genotypic landscapes of selected driver mutations with various degrees of ruggedness.

It was suggested that the ruggedness of landscapes determines the repeatability and predictability of adaptation[13]. Importantly, the ability to predict cancer progression using genomic data is an important goal in precision cancer medicine. Although we show that when the dynamics of the changing TME optimum is known/predetermined cancer adaptations may be predictable, previous studies that failed to consider a changing TME can significantly undermine the un-predictability of cancer evolution. Our results show that changing TMEs of any kind with Fisher's phenotypic fitness landscape can generate Sewall Wright's genotypic landscapes of selected driver mutations with considerable sign epistasis and ruggedness, which makes predicting cancer adaptation from genotypes alone in a changing TME challenging[14]. It is remarkable that simple phenotypic landscapes such as Fisher's can generate genotypic landscapes with such complexity[15]. Recent studies suggest even without a changing environment Fisher's phenotypic landscape can already generate complex genotypic landscapes with significant ruggedness[13,16]. However, under the classical assumptions, e.g., strong selection weak mutation-SSWM (per generation mutation is far less than 1 and the fate of a mutation can be determined), it may be possible to make predictions about the underlying adaptive evolution and give analytical results[17-22], which unfortunately does not apply to cancer evolution.

The requirement to track and sample the genotypes and phenotypes of cancer cell populations under selection in a time-dependent manner further makes predicting cancer evolution challenging[23], as these data are generally not accessible in clinical settings. Moreover, adaptive cancer evolution in individuals may have several evolutionary tempos and modes mixed across cancer types in humans, which can be further complicated by the heterogeneity of the TME and its varying optimum. On one hand, cancer cells may frequently go to extinction due to strong stabilizing selection from the normal TME. On the other hand, an extremely slow-changing or constant TME may lead, in effect, to limited cancer cell adaptation in which neutral/nearly neutral mutations accumulate, as might be the case for cancers that apparently carry no or few classical driver mutations. Of course, the TME may determine the phenotypic effect of a mutation as well as determine the fitness landscape, so a changing TME might alter both a phenotype and its selective advantage. Furthermore, cancer growth and progression (e.g. acquisition of new mutations) will also affect the TME; and the cancer itself may create or modify a TME optimum leading to non-cell-autonomous cancer evolution. Therefore, quantitative understanding of these basic components and phenotypic optimum change dynamics in individual patients is crucial for developing future predictive cancer medicine. For instance, we show that a cycling TME may be particularly capable of promoting cancer adaptation (see Supplementary Movies S16-S17 for its unusual long-term sub-clonal dynamics and a large number of recorded adaptive steps), while stochasticity and fast changes in the TME optimum may act against cancer adaptation and cause extinction. TME dynamics need to be considered alongside other factors, such as phenotypic and genetic factors, in studies that aim to provide a complete picture of the underlying cancer evolutionary dynamics. Finally, cancer research must consider the significance of a changing tumour microenvironment in cancer progression and treatment.

#### Supplementary Note 7

#### Changing phenotypic optimum and spatial constraints affect cancer phylogenies and adaptation

Here we further describe the characteristics of phylogeny shape from the evolving cancer under randomly or cyclically changing phenotypic optimum. In randomly changing TMEs (Figure S25 **a-d**), the higher optimum change variance leads to shorter side branches indicating higher rates of stochastic death of these cancer cell lineages, which disfavours adaptive evolution, as evidenced by the lack of obvious asymmetric (imbalanced) trees. In a cycling TME (Figure S25e-h), a low peak of the optimum changing cycle (equation (13)) leads to a phylogeny showing recent clonal expansion (Figure S25**e**), whereas an intermediate amplitude promotes adaptive evolution (Figure S25**j**), as evident from the phylogenies' ladder-like and spindly tree topology (strongly imbalanced and asymmetric). The temporal signals in the phylogeny also reflect a cycling pattern, where long branches and ladder-like shorter branches appear in tandem (figure S25**f**).

We then quantify the adaptation and phylogeny shapes (using the normalised Sackin's index and the number of cherries) of the evolving cancer under the three different (directional, cyclic and random) phenotypic optimum change dynamics with different levels of spatial constraints. In all three types, increasing the changing speed of the phenotypic optimum reduces mean population fitness and selects for driver mutations with large selective advantage. However, stronger spatial constraints on population size (smaller space) further select for driver mutations with increased selective advantage in both directionally and randomly changing TMEs (Supplementary Figure S26a-c) but with decreased mean population fitness for all three types of TMEs (Supplementary Figure S26d-f). Complementarily, the two phylogeny shape measures also reveal the underlying evolutionary processes (Supplementary Figure S27). When the phenotypic optimum changes fast, the two measures of phylogeny shape suggest there is an increase in selection-induced phylogeny shape asymmetry at both global and local levels. However, lifting spatial constraints on population size (increasing space size) leads to increases in both the number of cherries (Supplementary Figure S27a-c) and normalised Sackin's index (Supplementary Figure S27d-f). Interestingly, when the space size has reached certain level in this strong selection regime, space increase only leads to a marginal increase in the number of cherries indicating recent asymmetry at each sampling point is less affected (Supplementary Figure S27a-c). All these results suggest the spatial effect on tree asymmetry and cancer adaptation is more profound when the phenotypic optimum changes fast.
