## Supplementary Figure S1 for "Why is cancer not more common? A changing microenvironment may help to explain why, and suggests strategies for anti-cancer therapy"

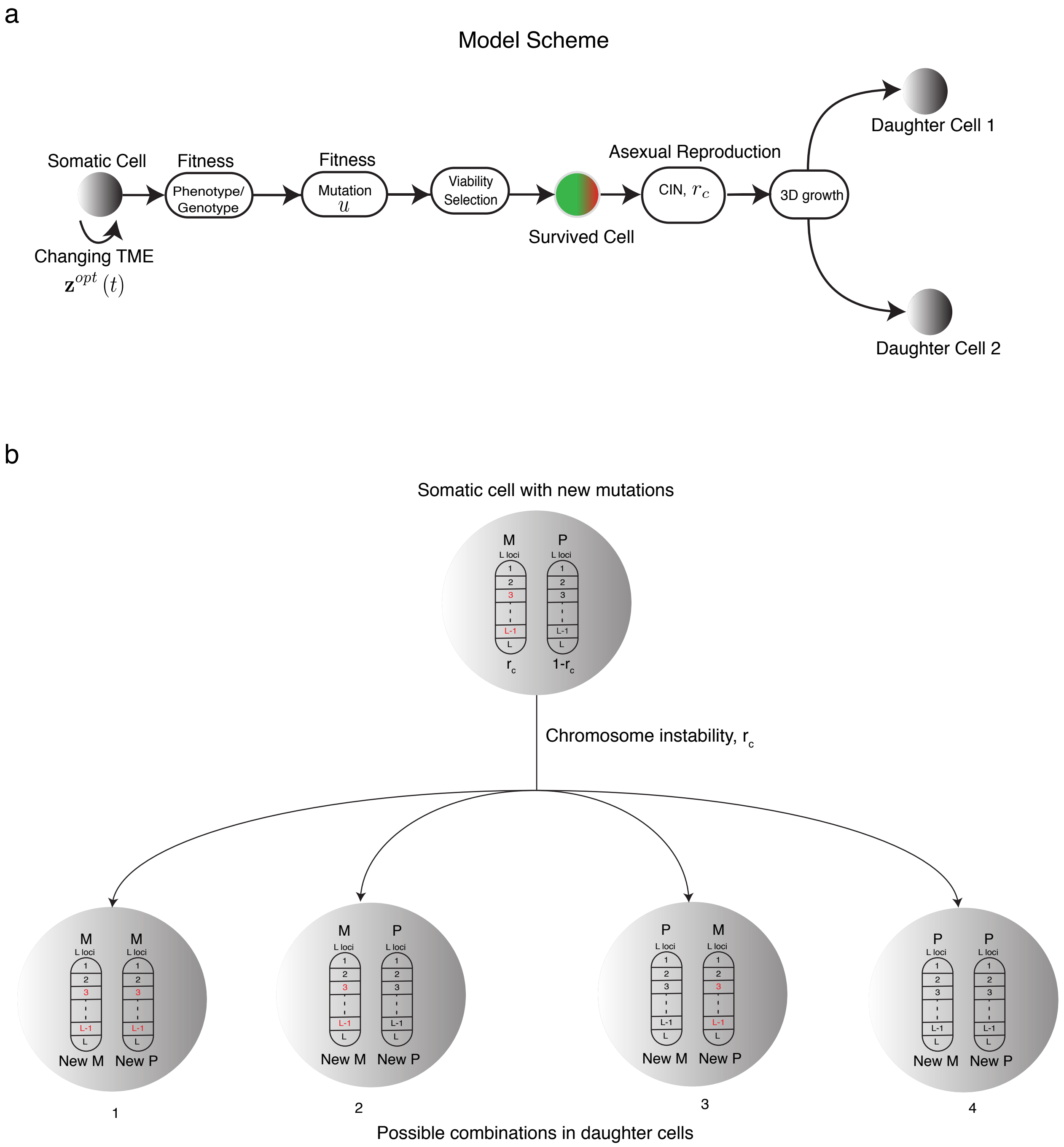


Symbols and Abbreviations:

| | |
|---|---|
| $\mathbf{z}^{opt}(t)$ | Tumour microenvironment (TME) changing dynamics |
| $u$ | Mutation rate |
| $r_c$ | Chromosome instability (CIN) rate |
| M | Maternal chromosome |
| P | Paternal chromosome |
