## Supplementary figures and images for "Why is cancer not more common? A changing microenvironment may help to explain why, and suggests strategies for anti-cancer therapy"

### Supplementary Figure S2

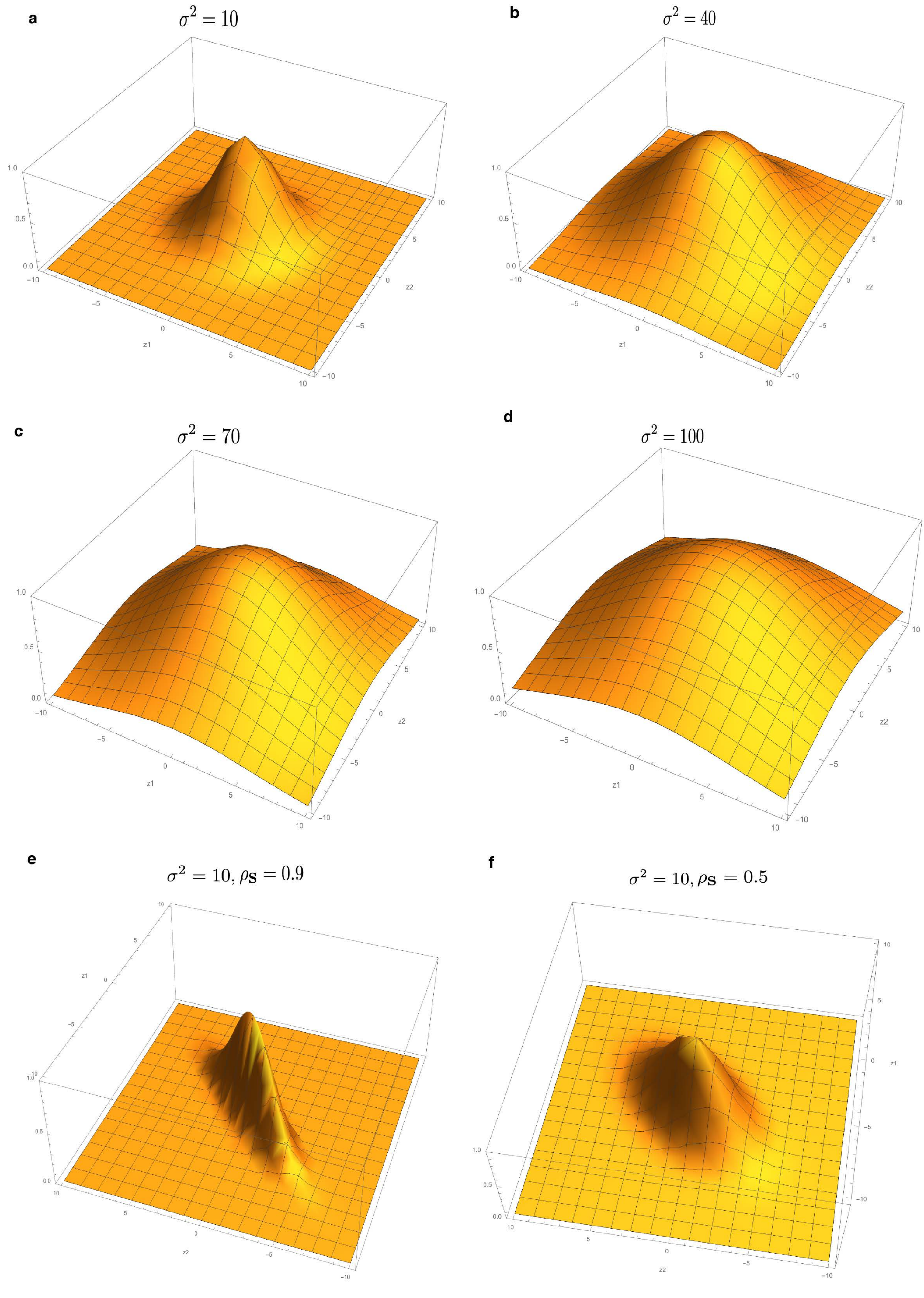

### Supplementary Figure S3

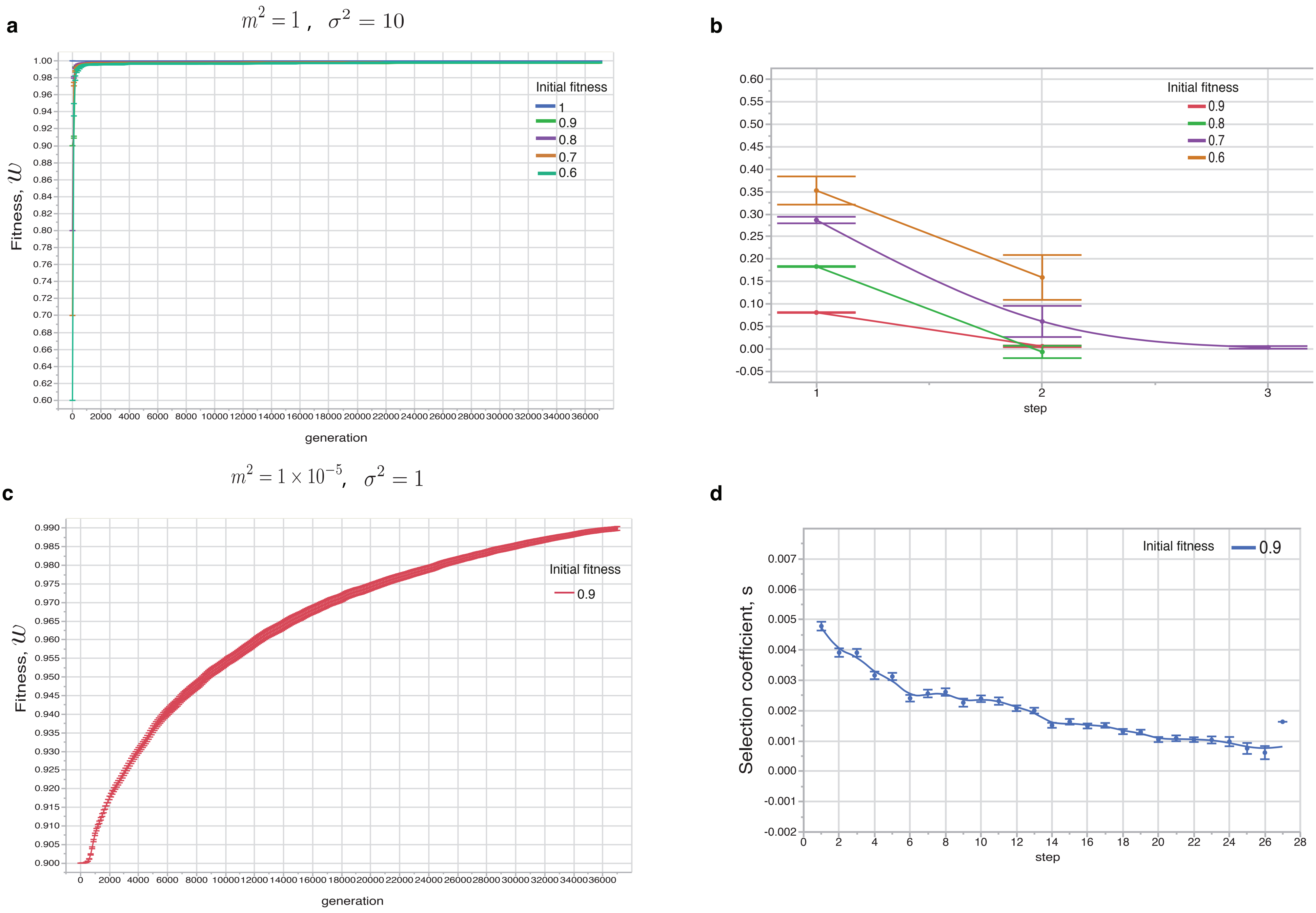

### Supplementary Figure S4

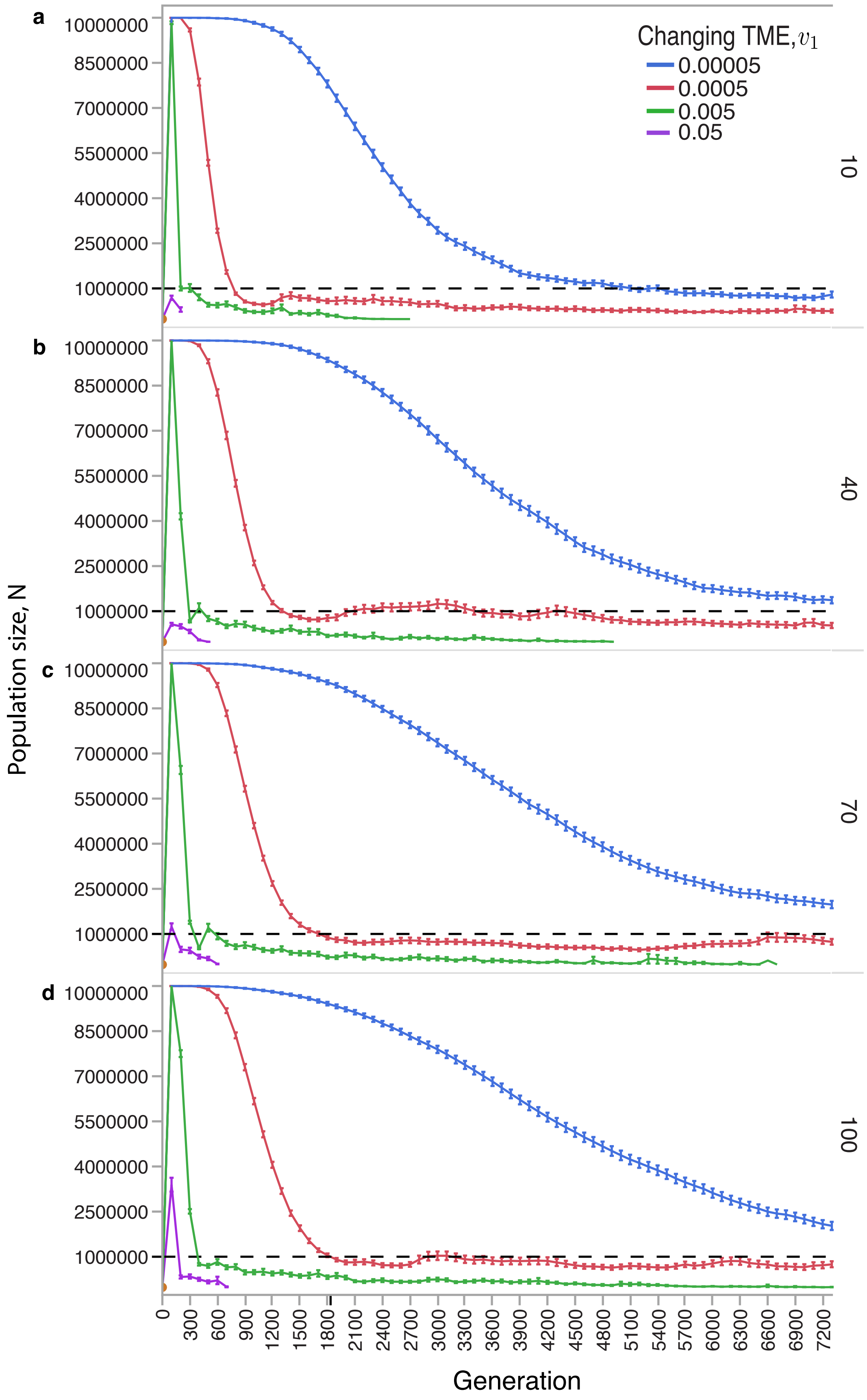

### Supplementary Figure S5

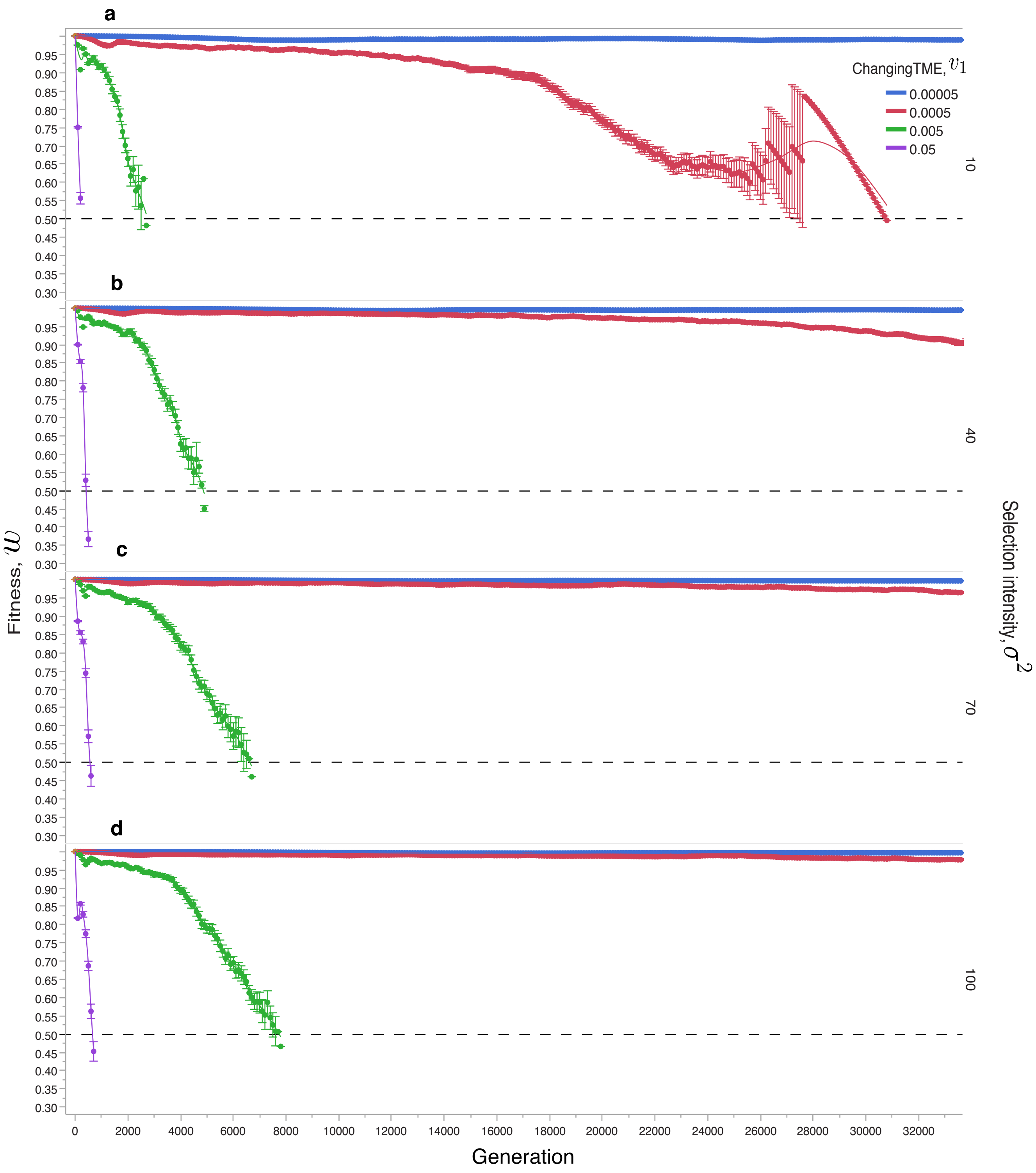

### Supplementary Figure S6

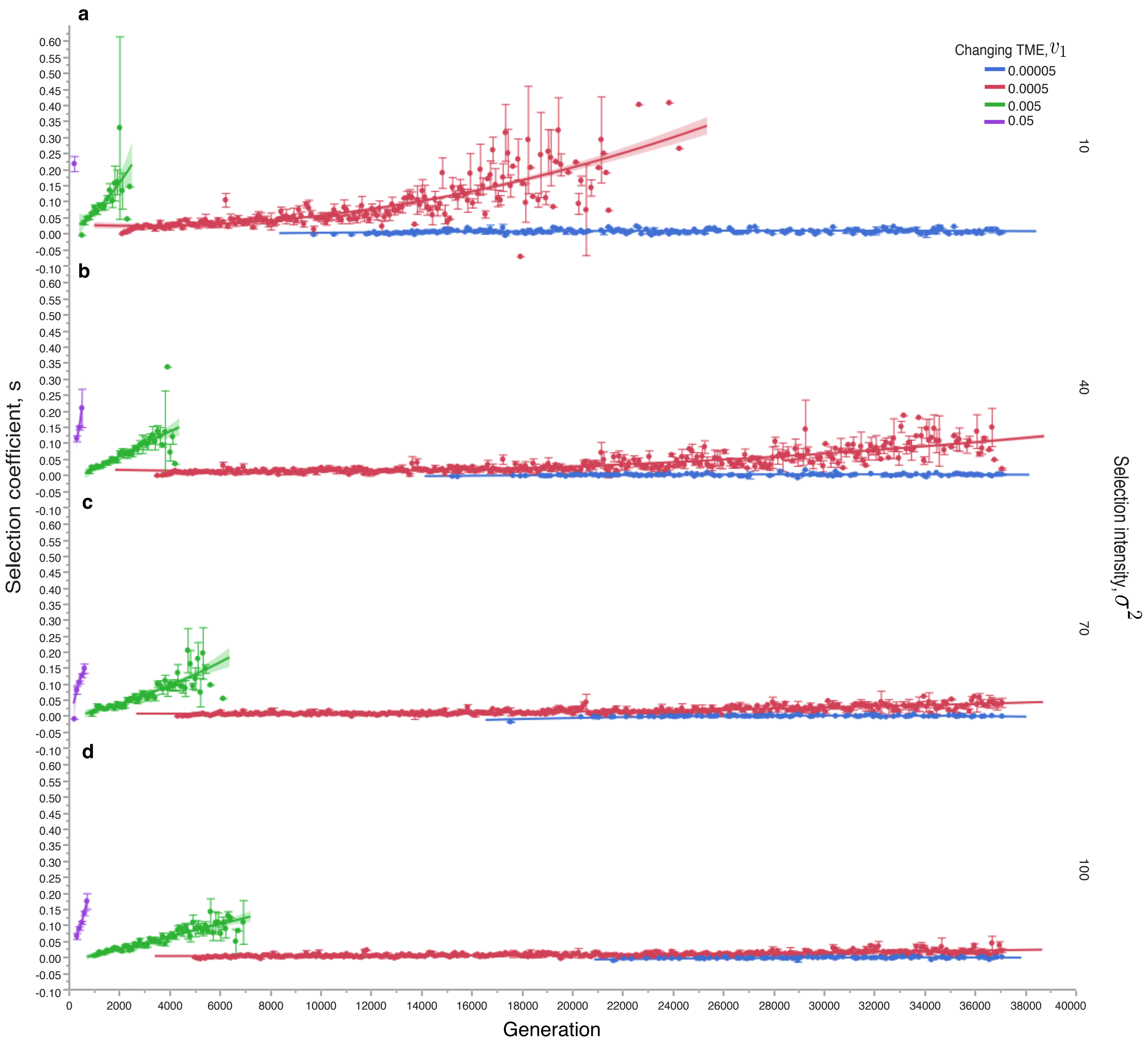

### Supplementary Figure S7

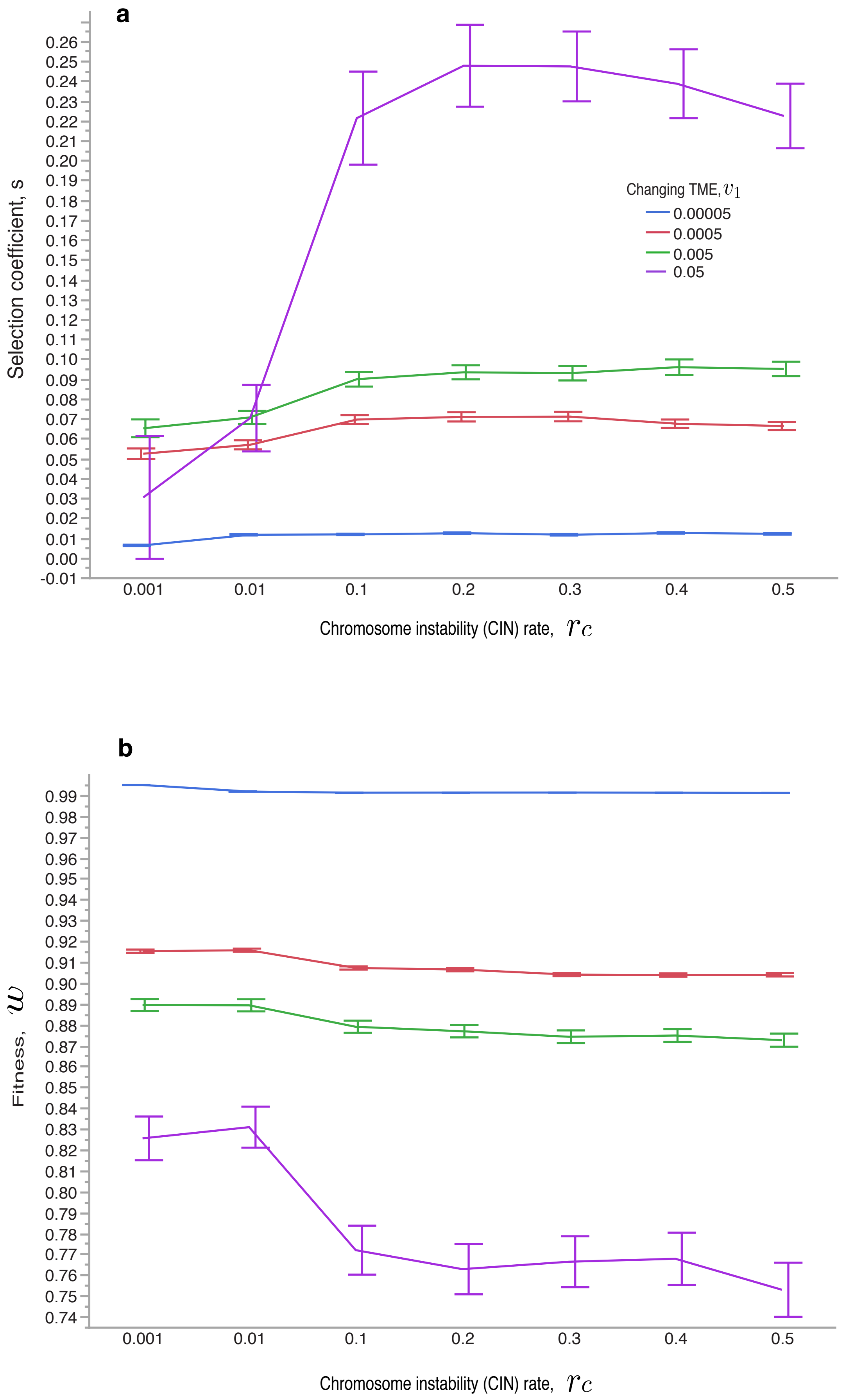

### Supplementary Figure S8

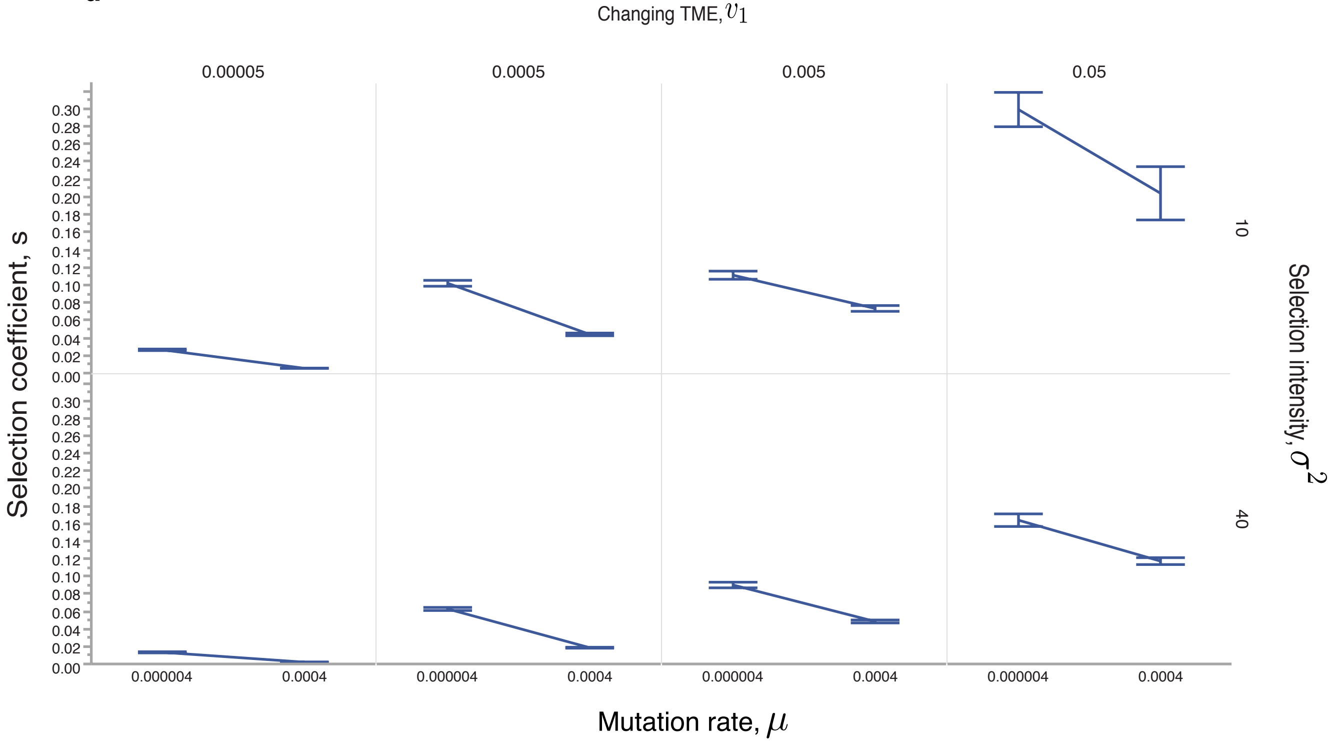

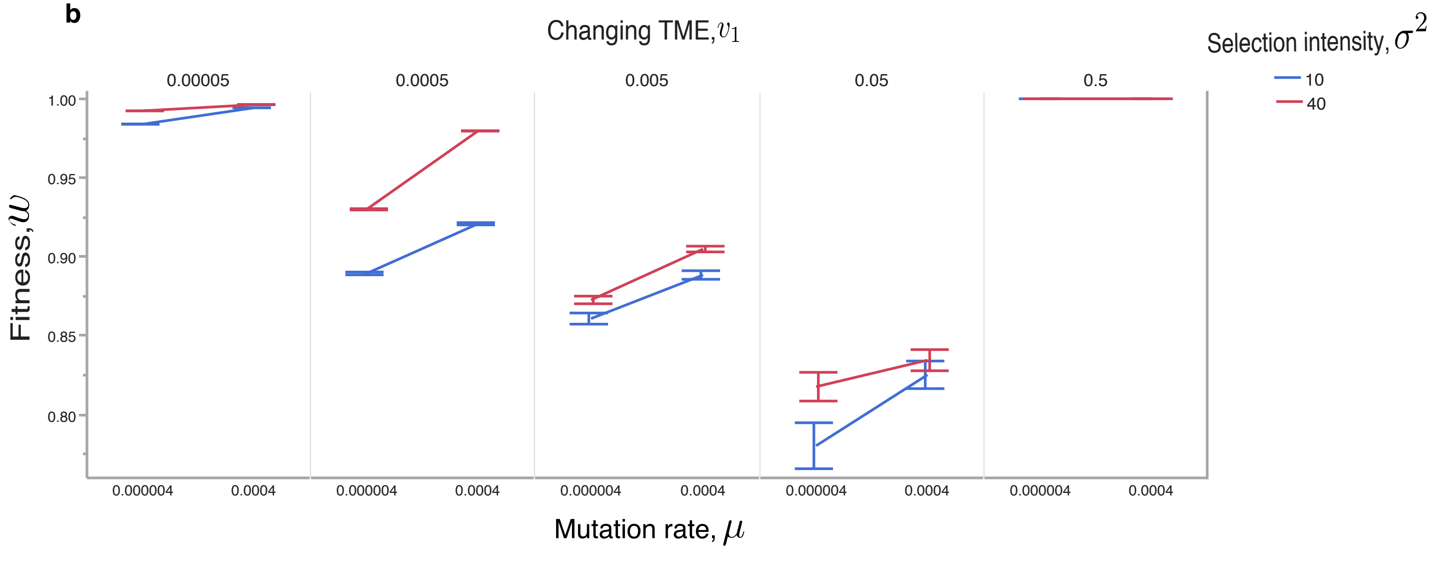

### Supplementary Figure S9

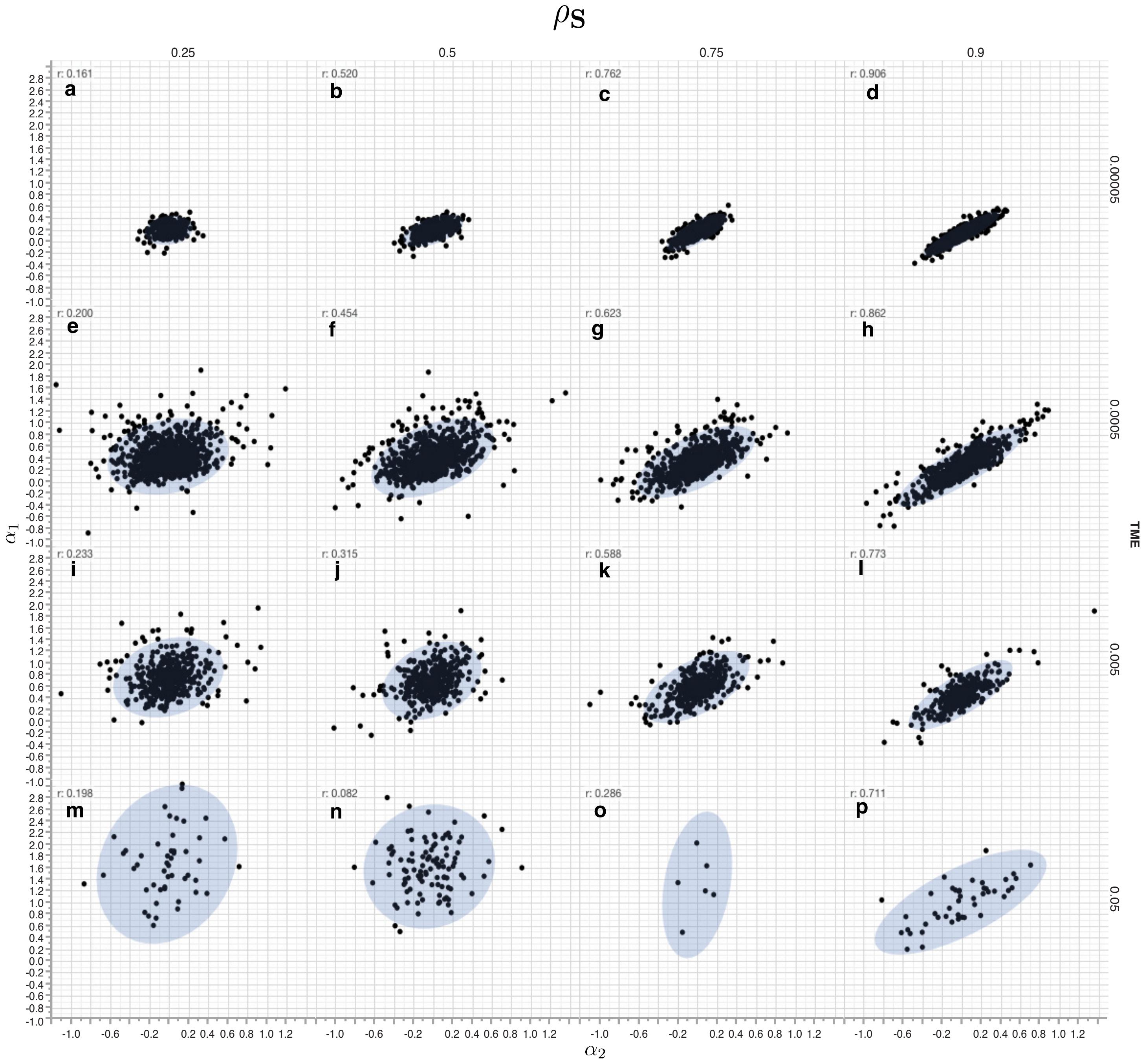

### Supplementary Figure S10

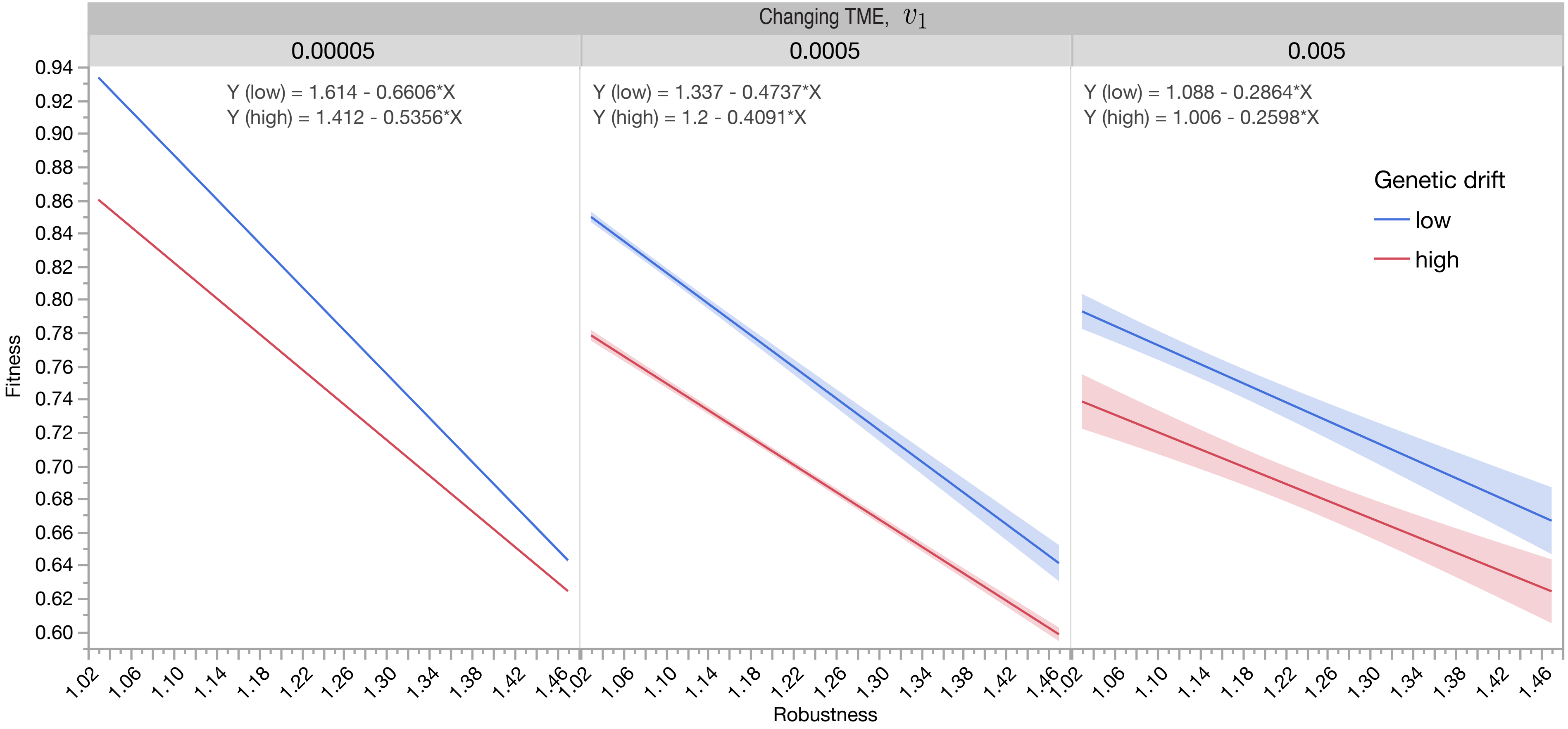

### Supplementary Figure S11

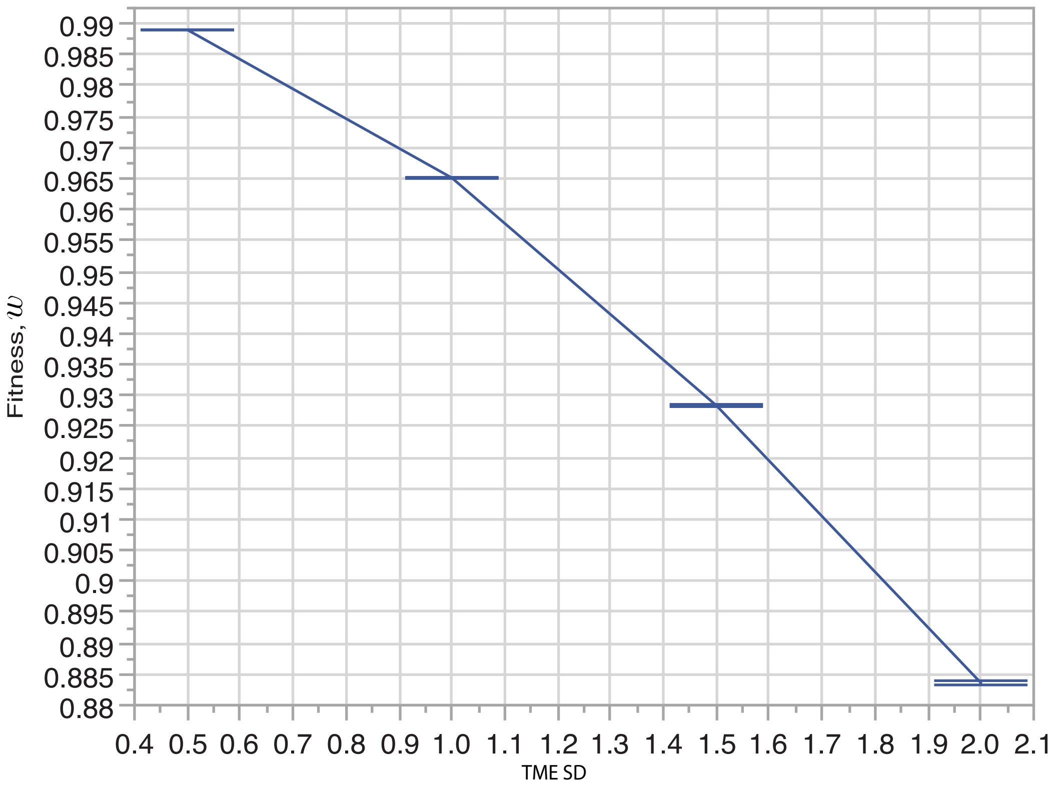

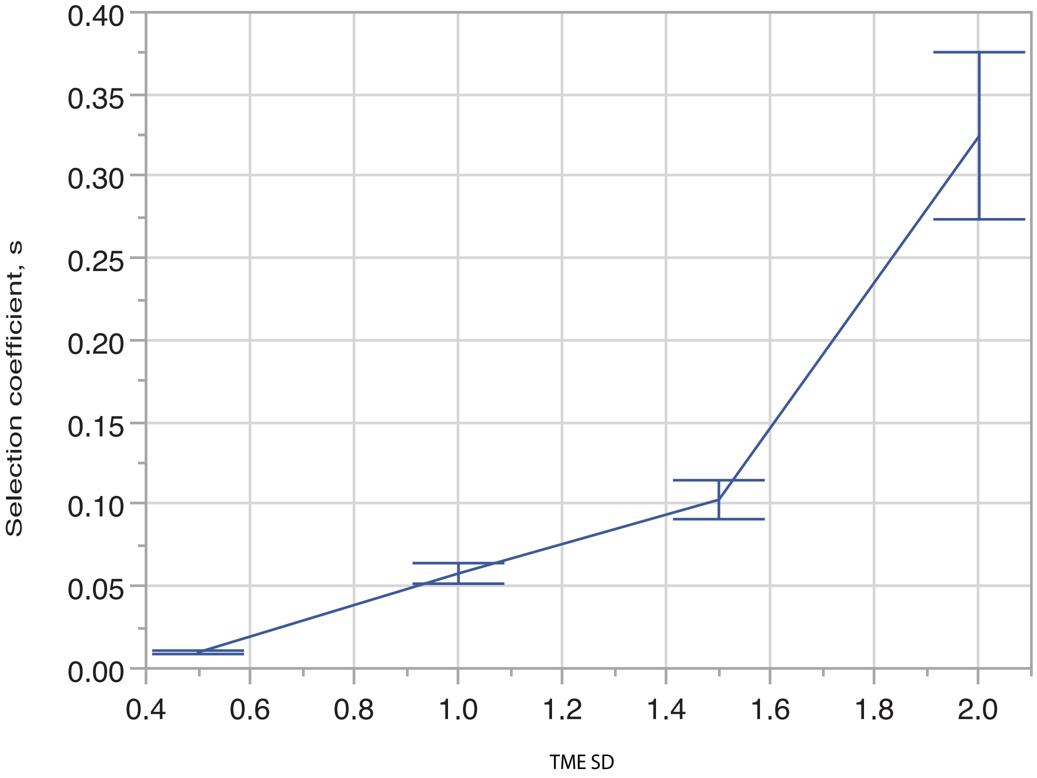

### Supplementary Figure S12

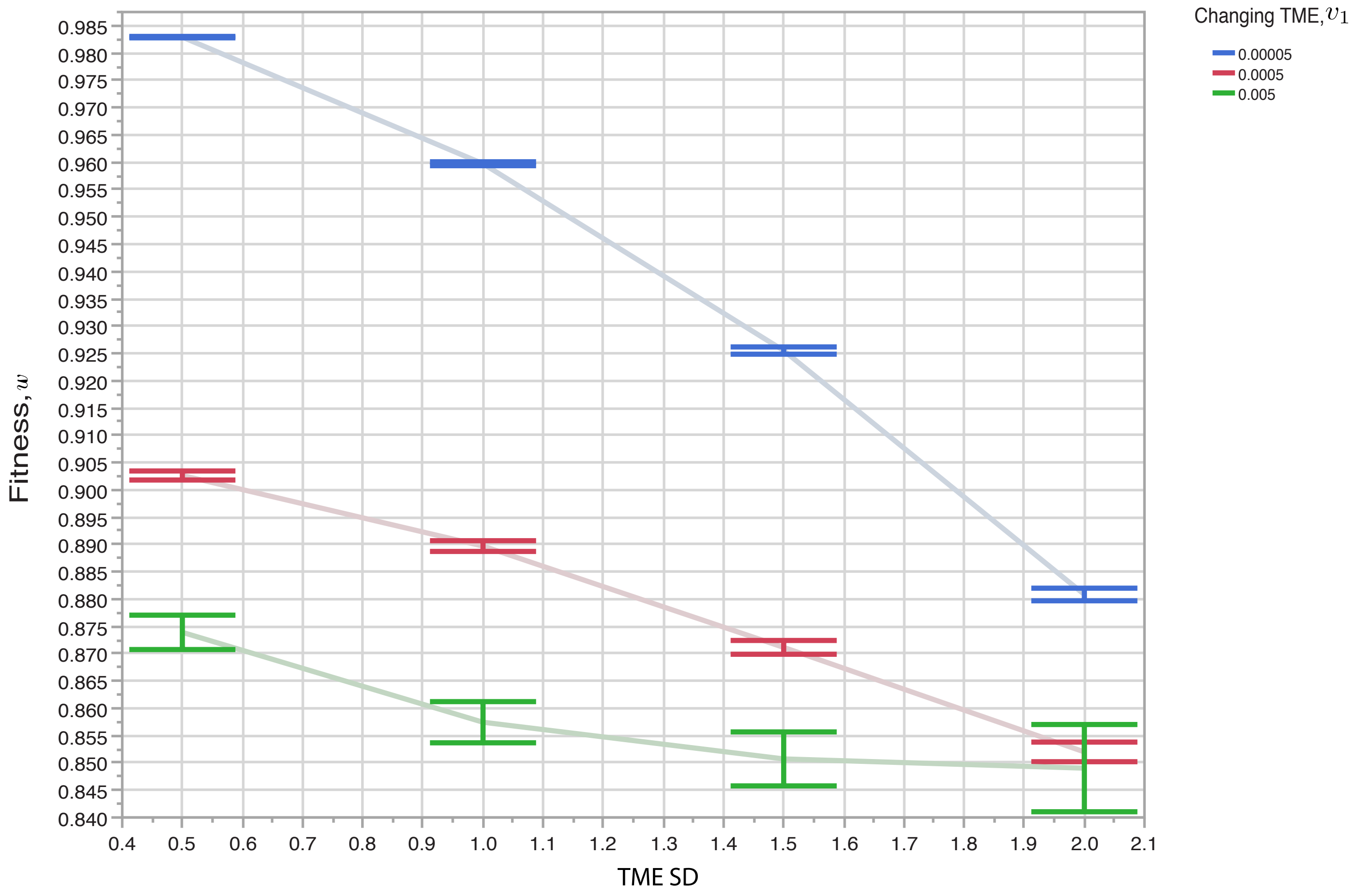

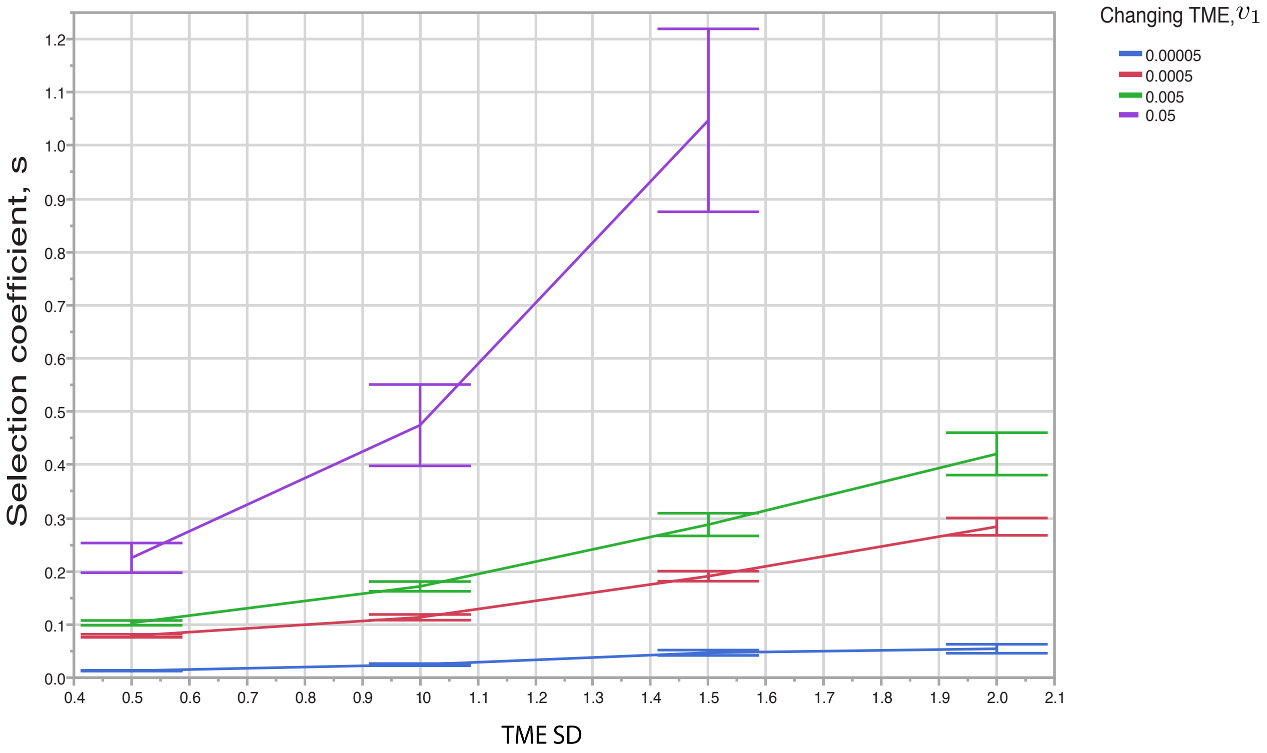

### Supplementary Figure S13

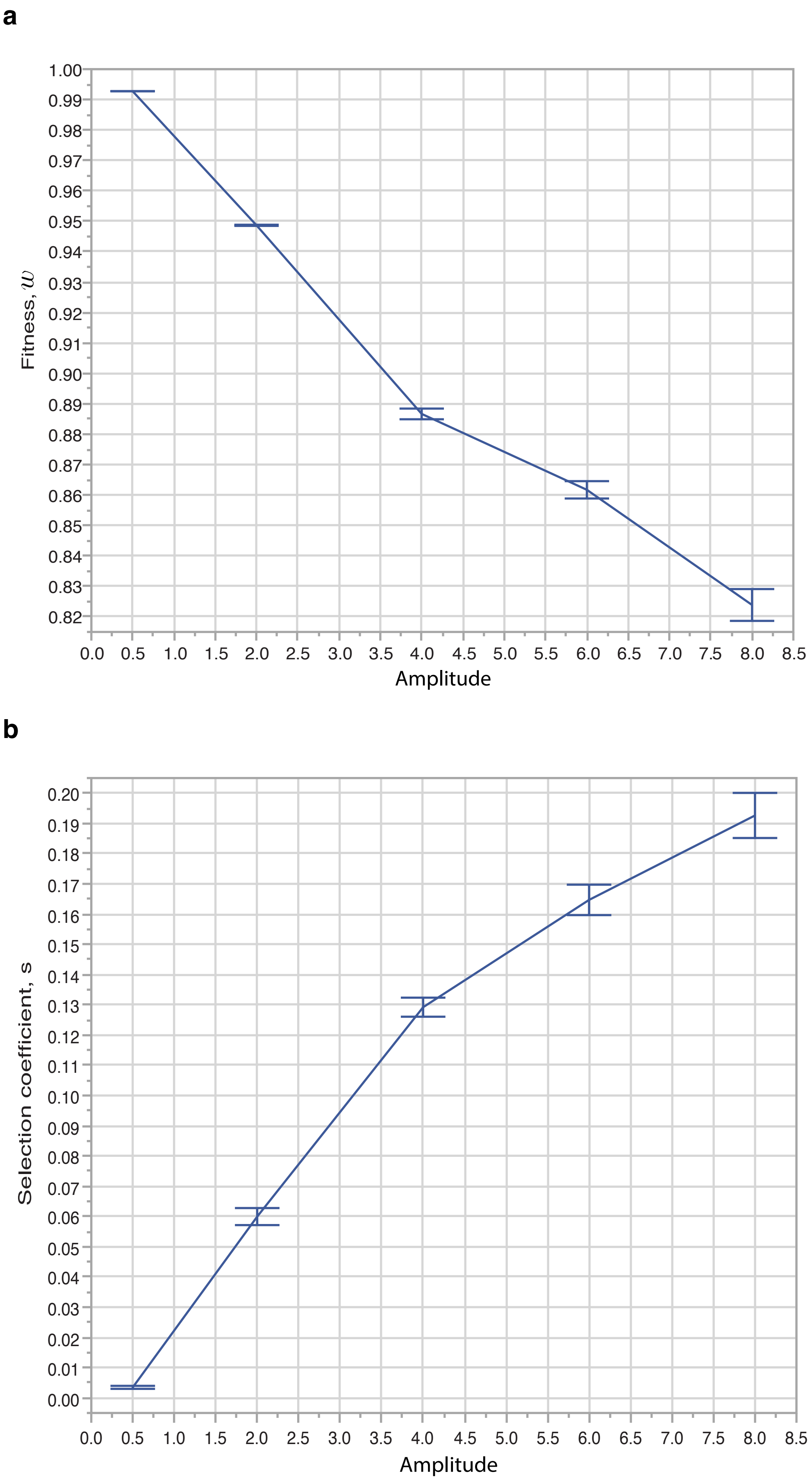

### Supplementary Figure S14

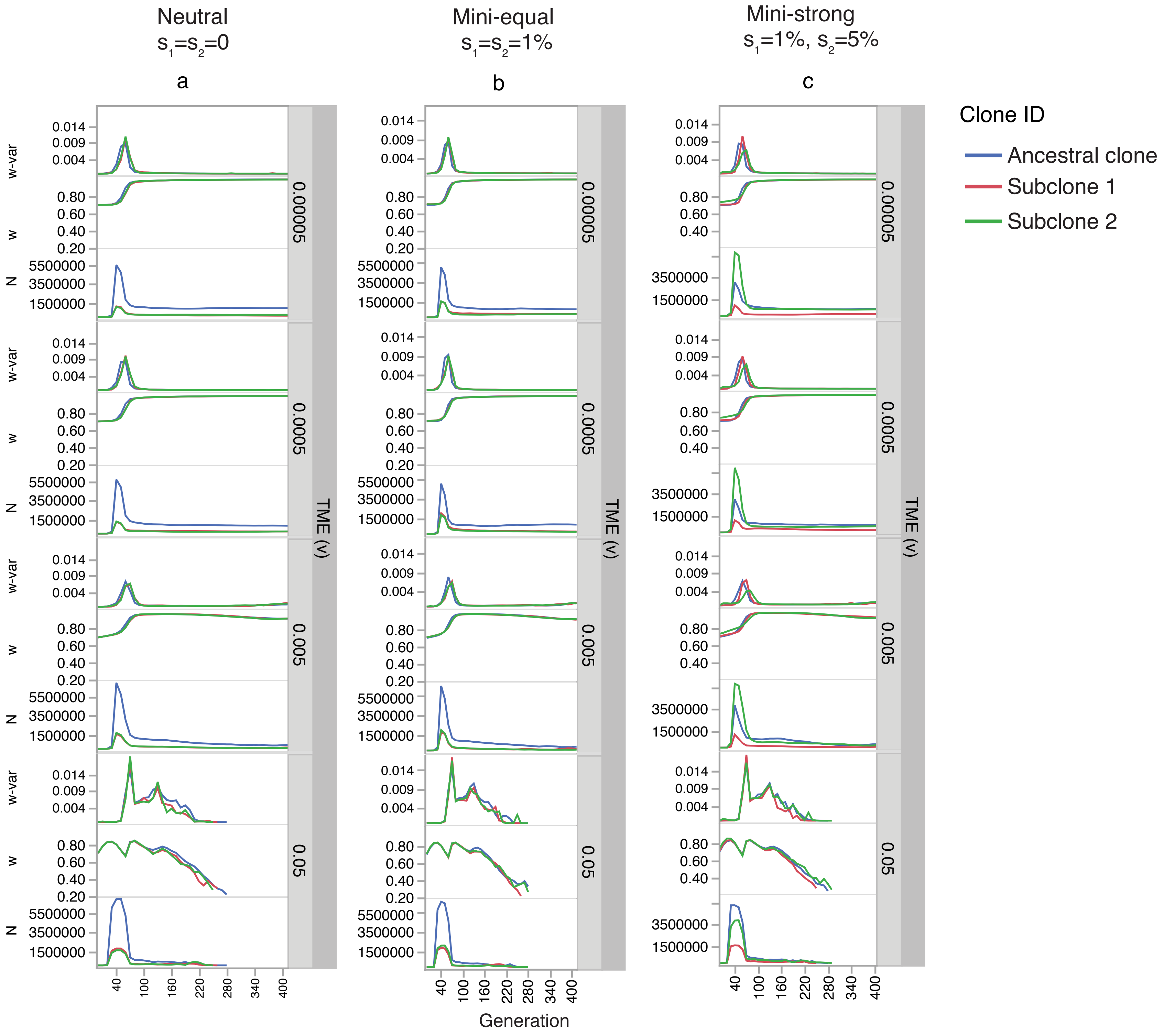

### Supplementary Figure S15

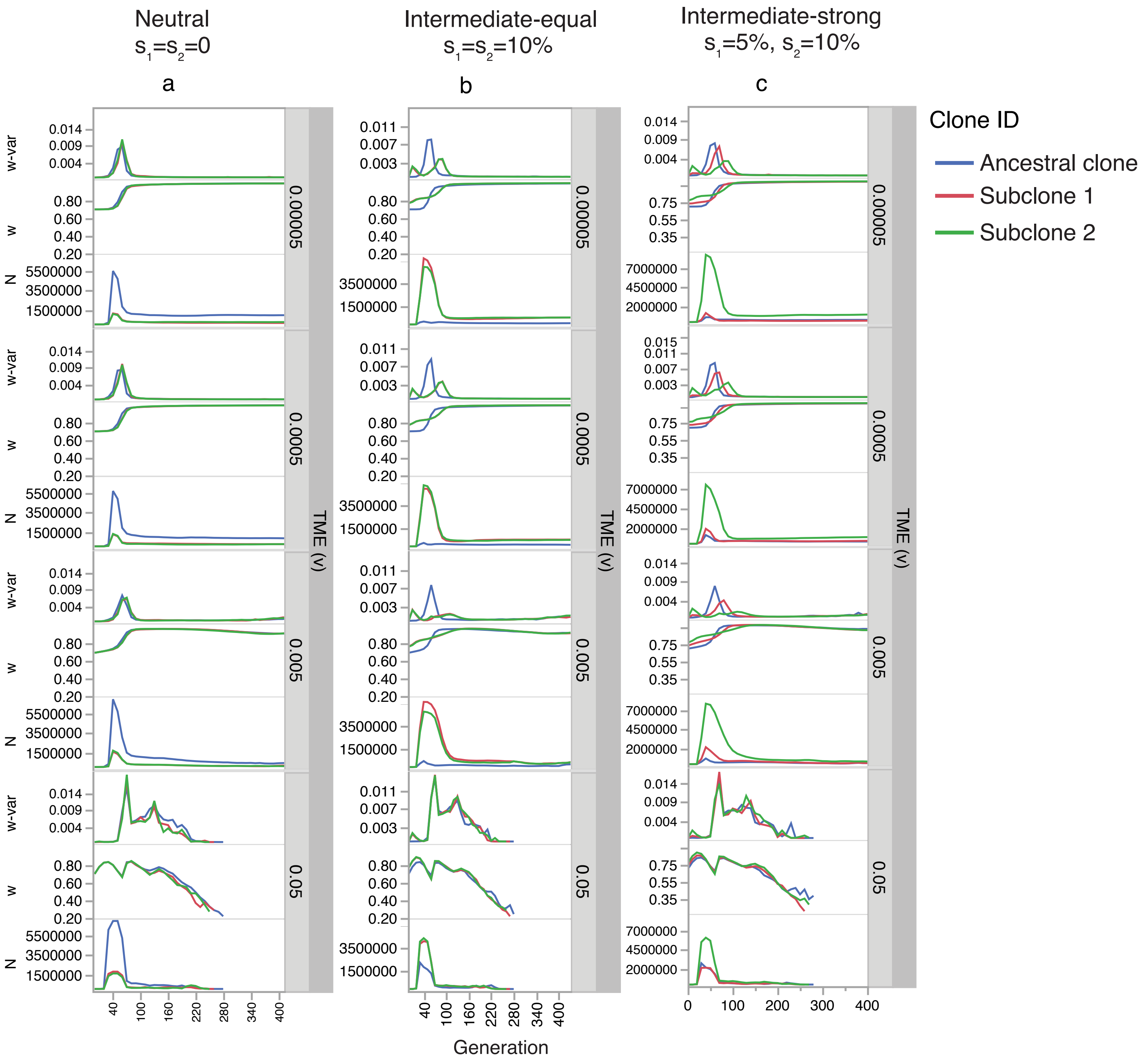

### Supplementary Figure S16

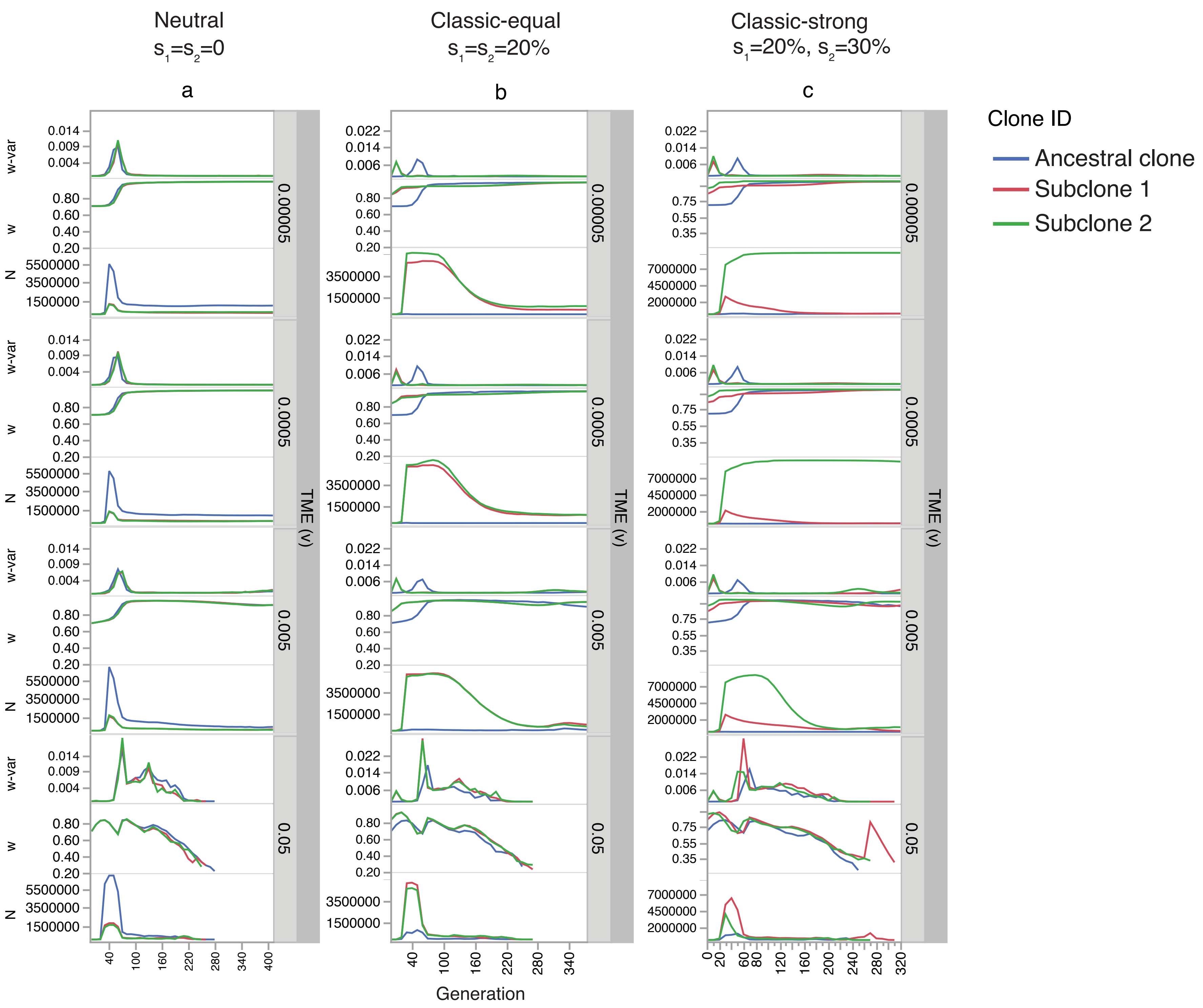

### Supplementary Figure S17

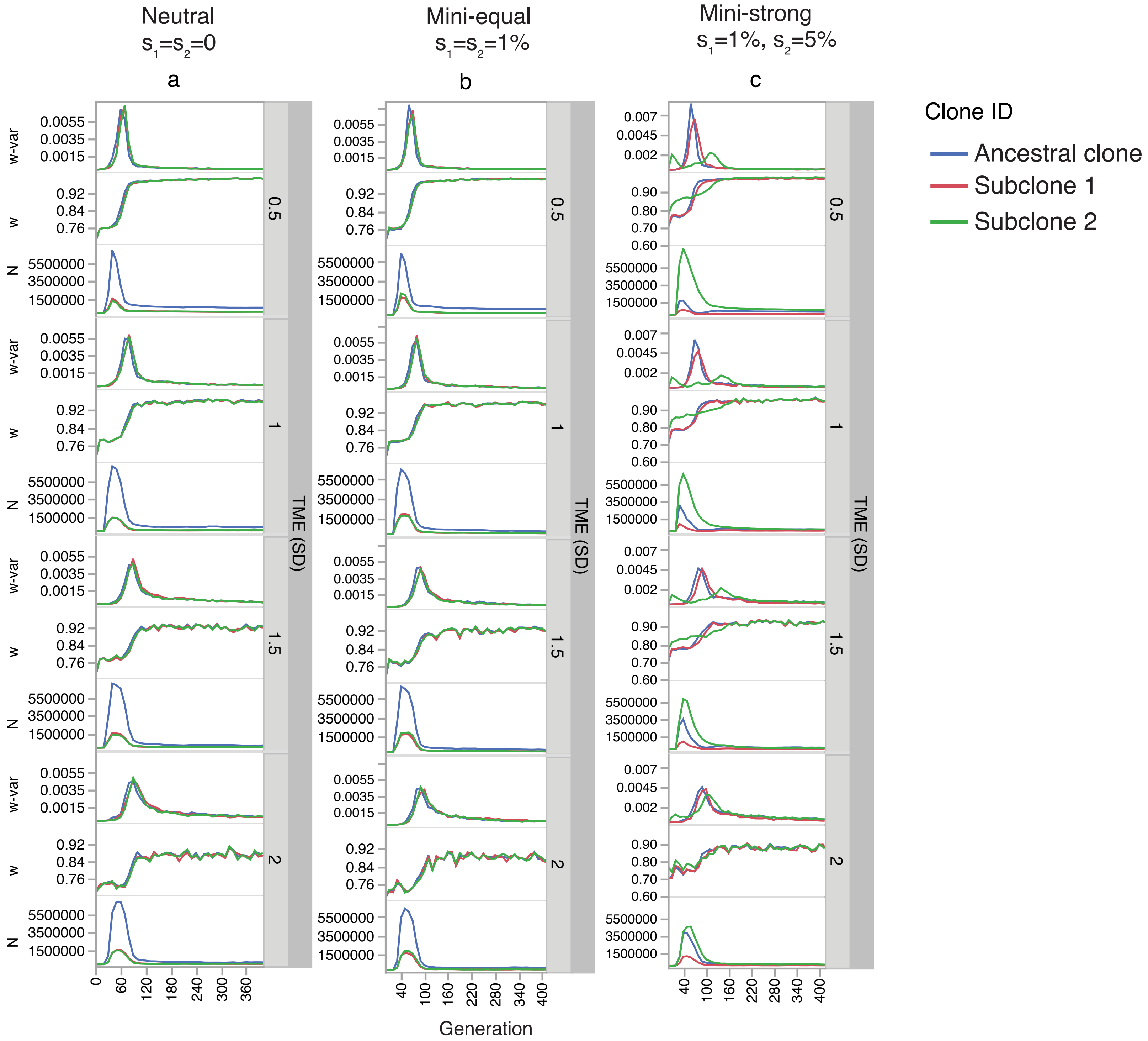

### Supplementary Figure S18

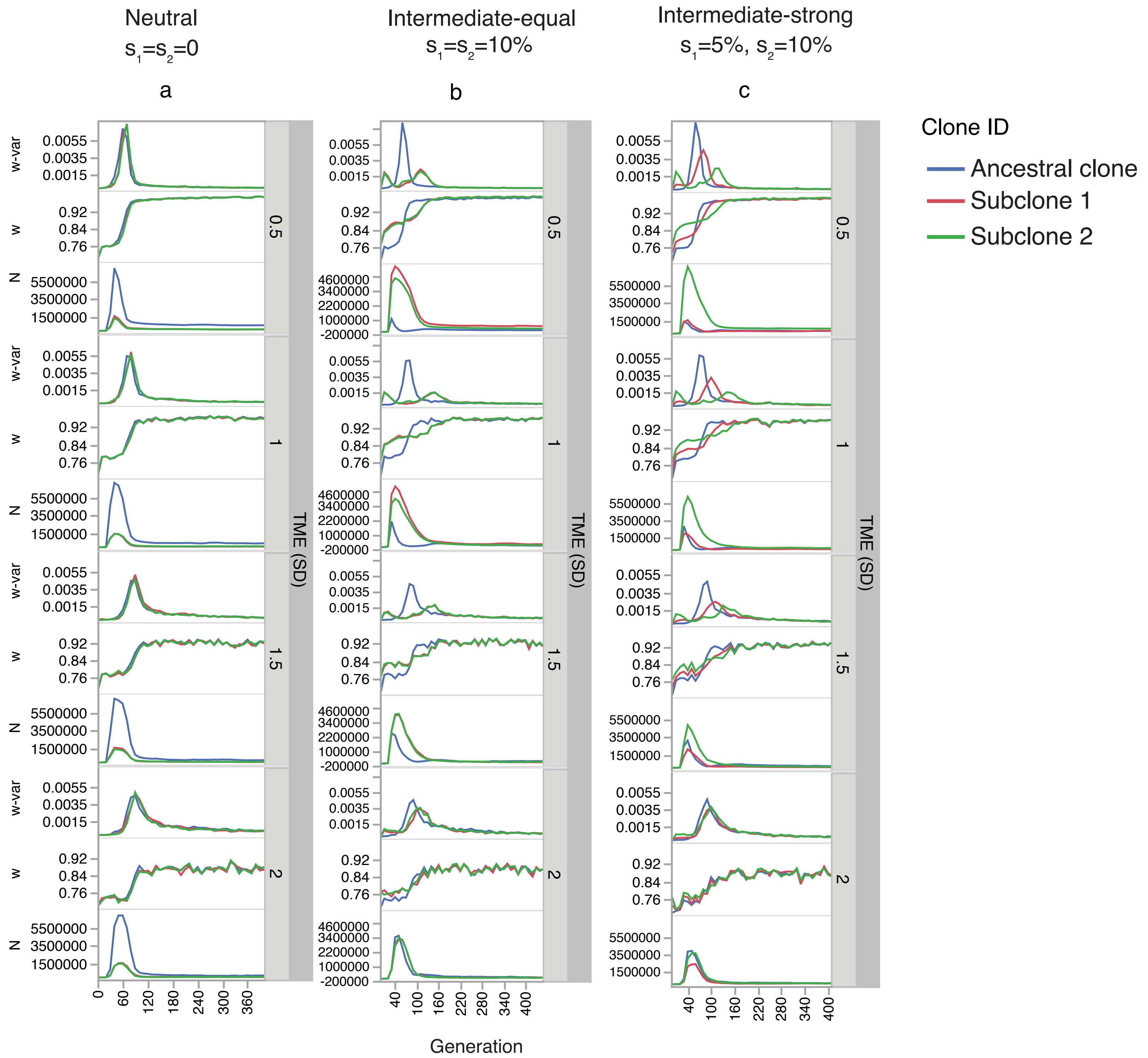

### Supplementary Figure S19

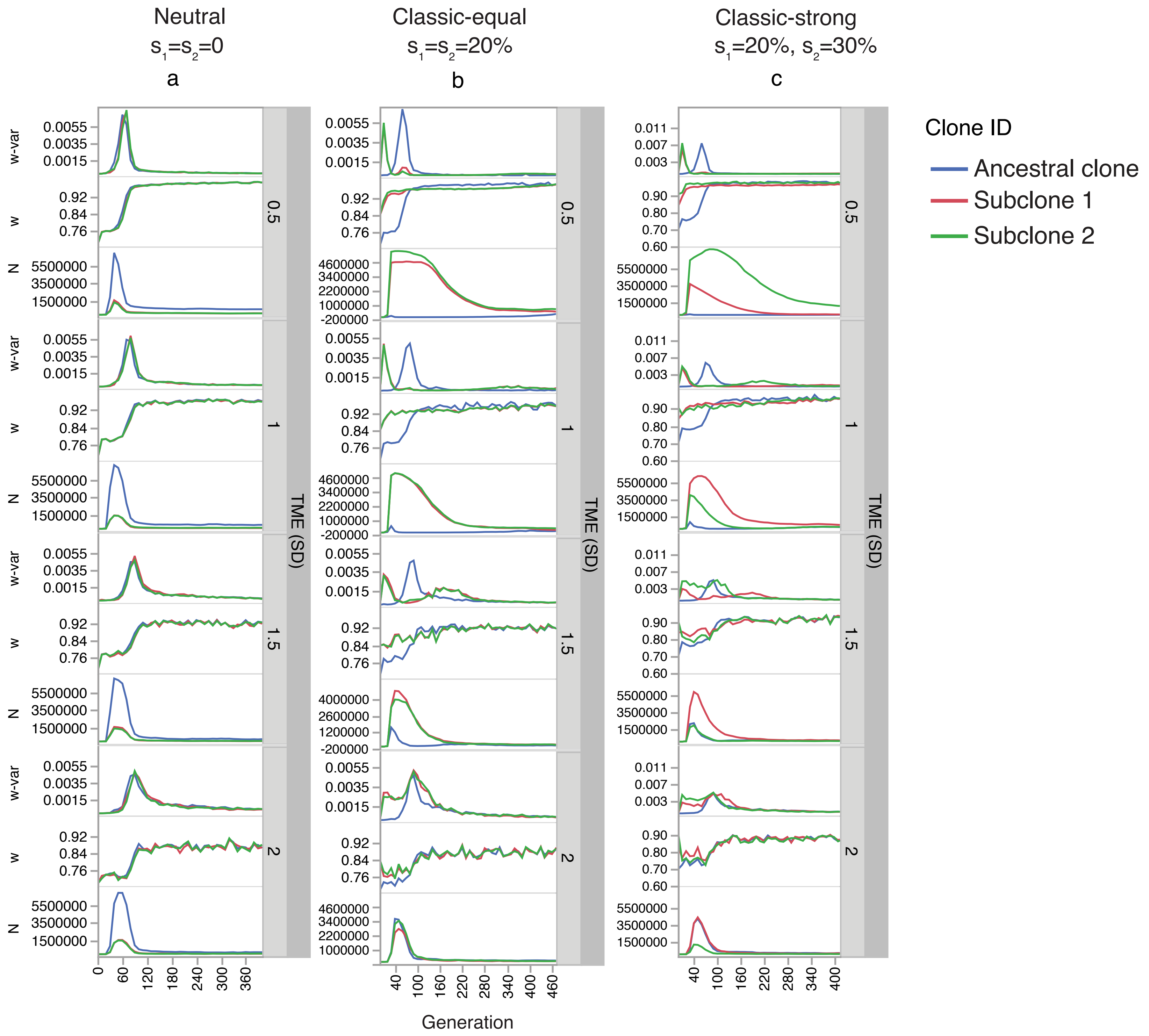

### Supplementary Figure S20

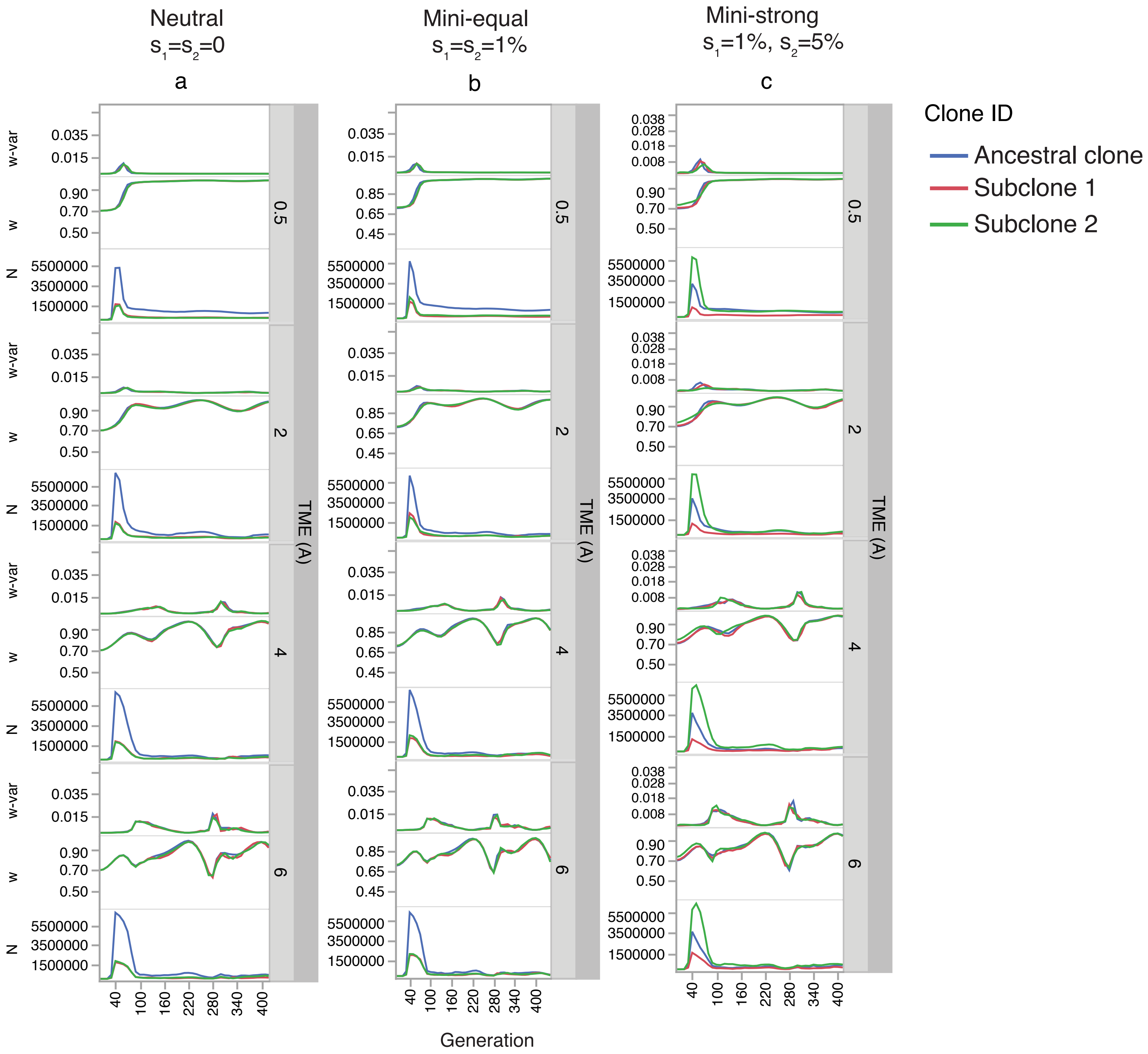

### Supplementary Figure S21

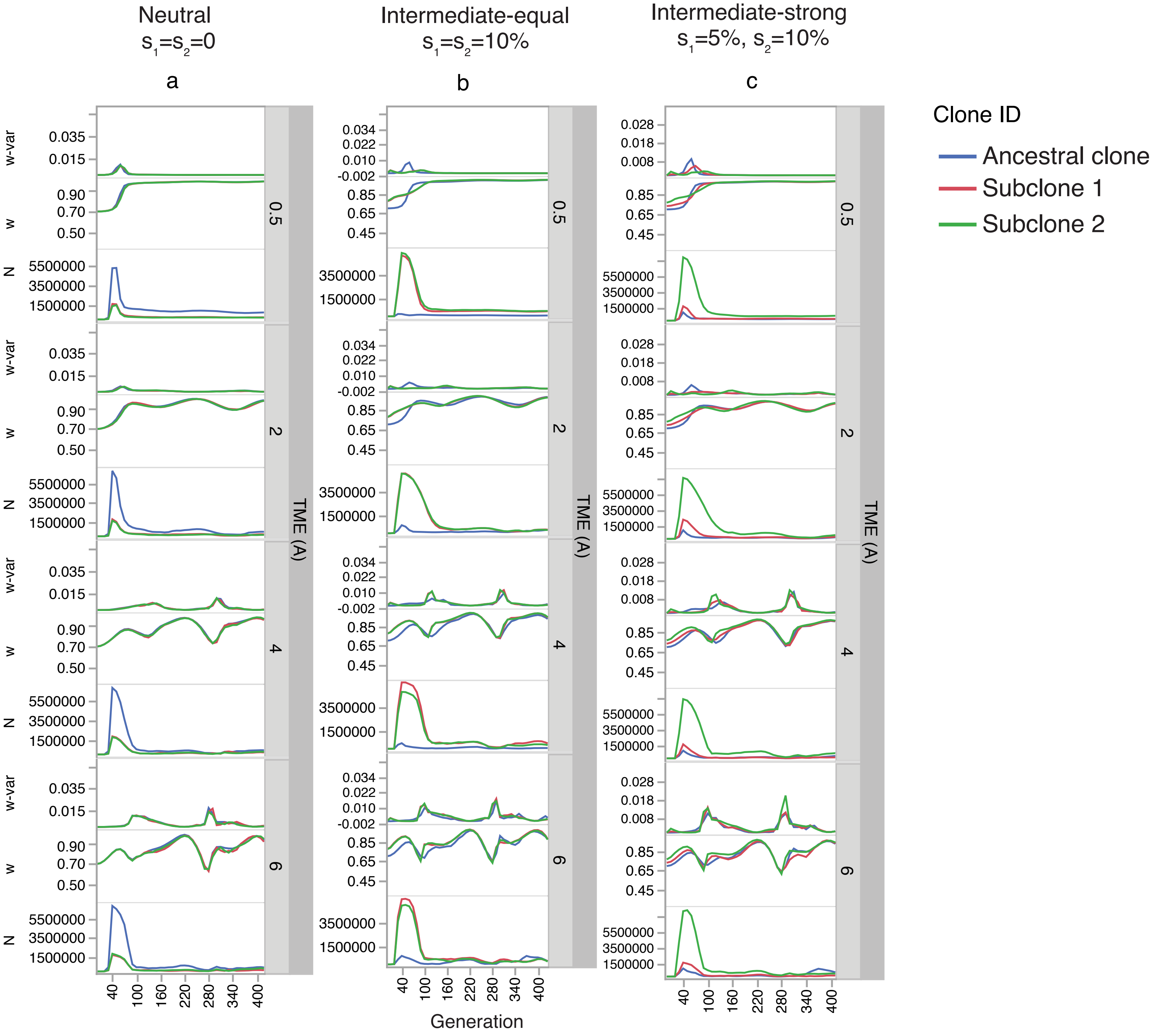

### Supplementary Figure S22

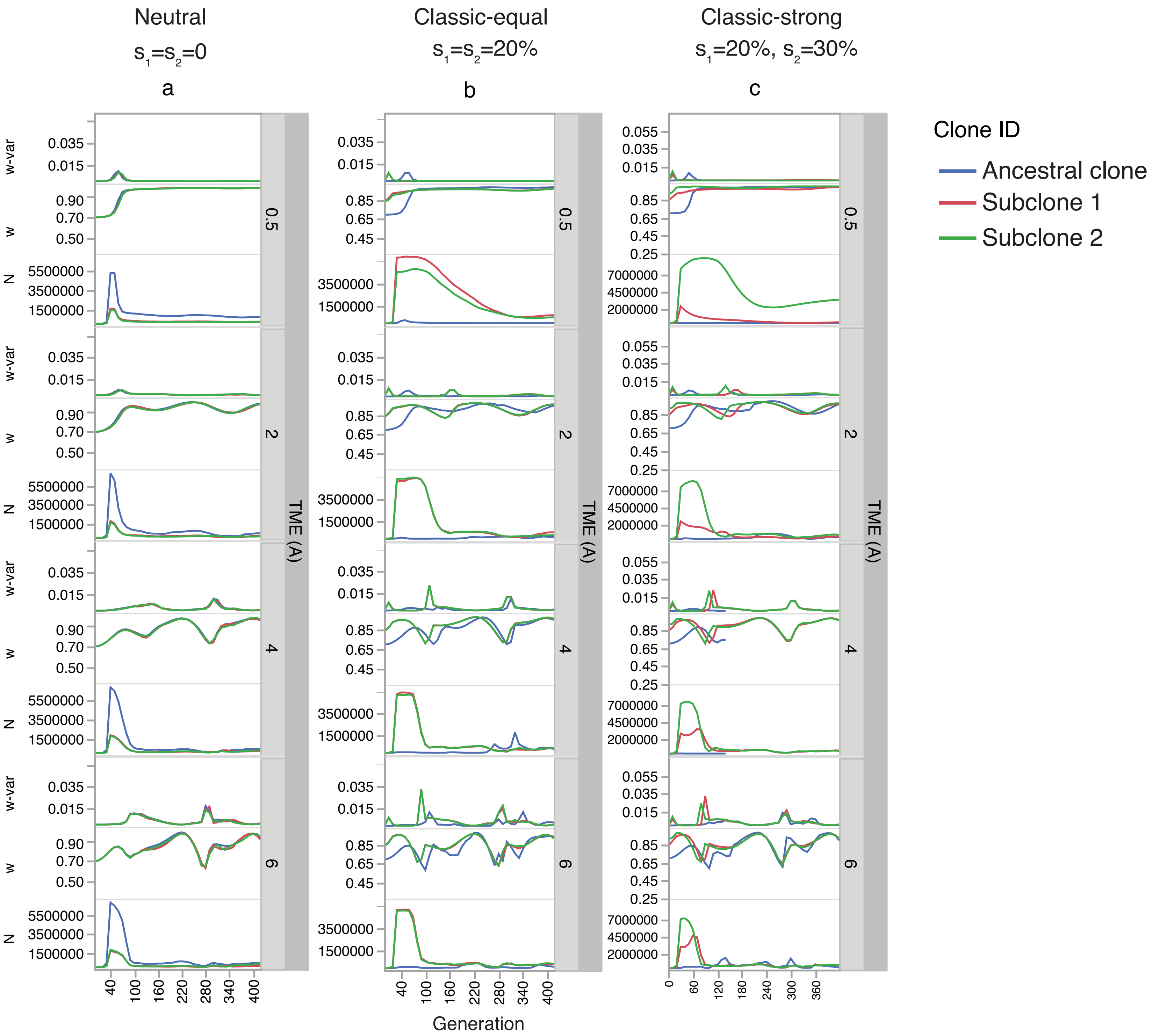

### Supplementary Figure S23

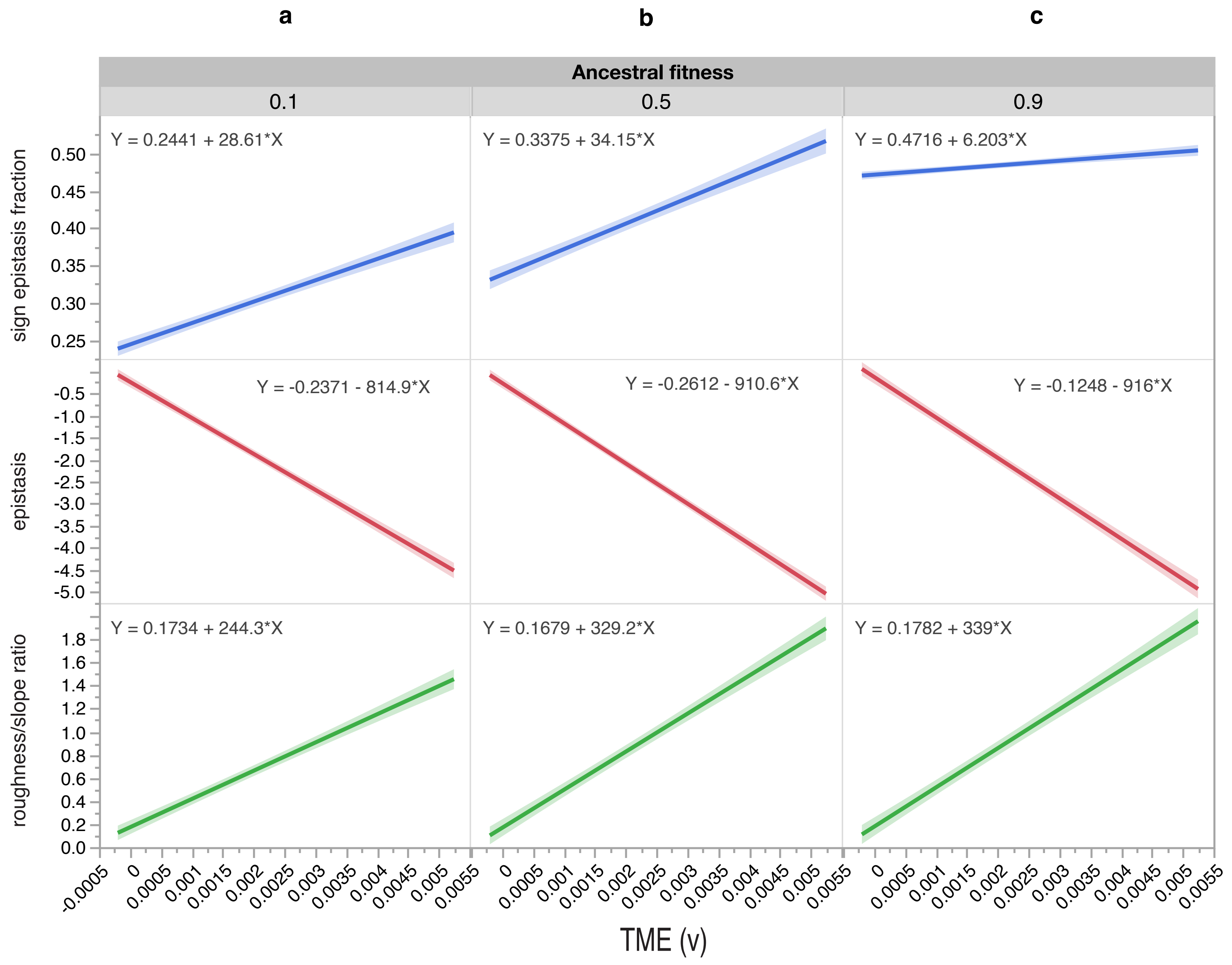

### Supplementary Figure S24

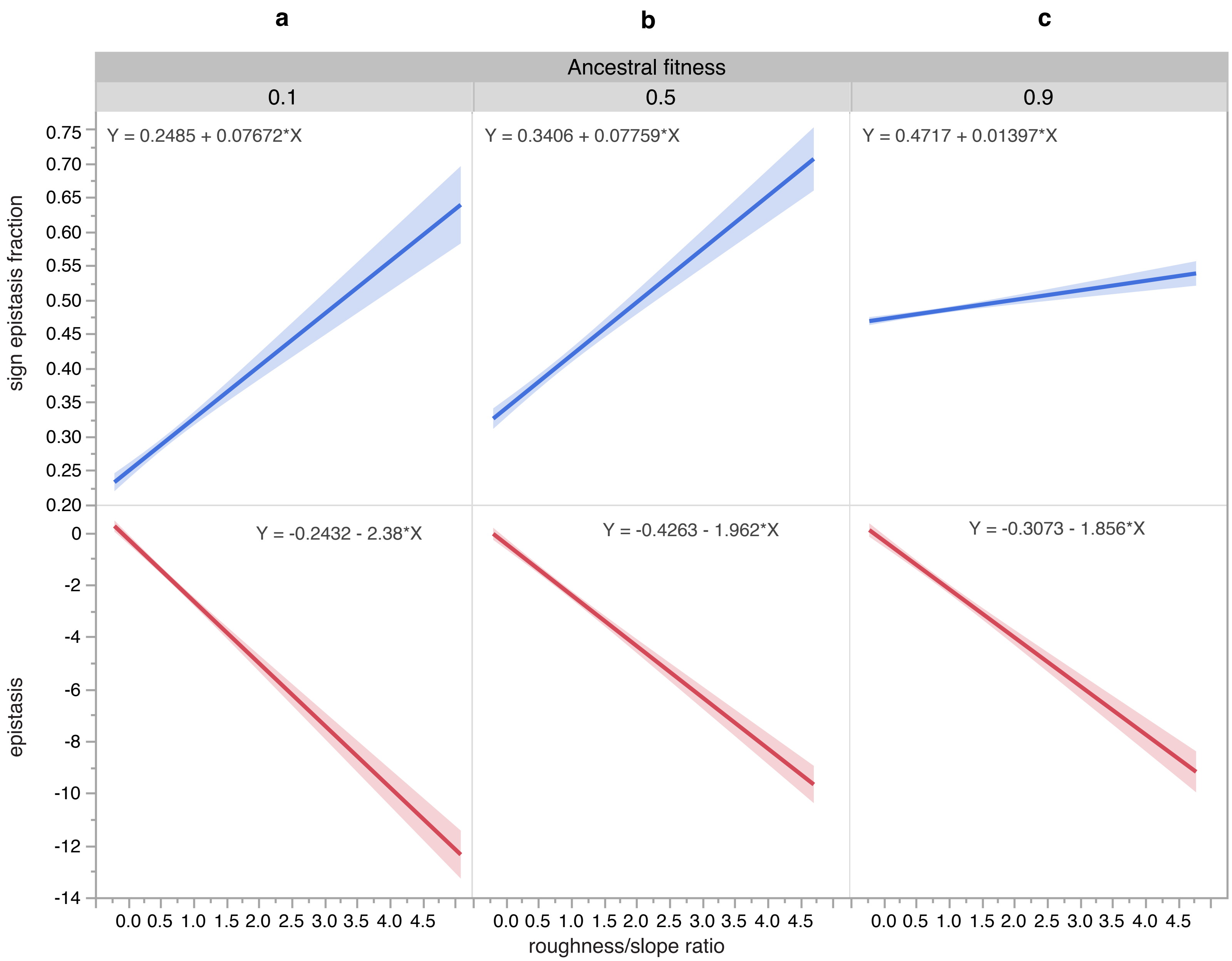

### Supplementary Figure S25

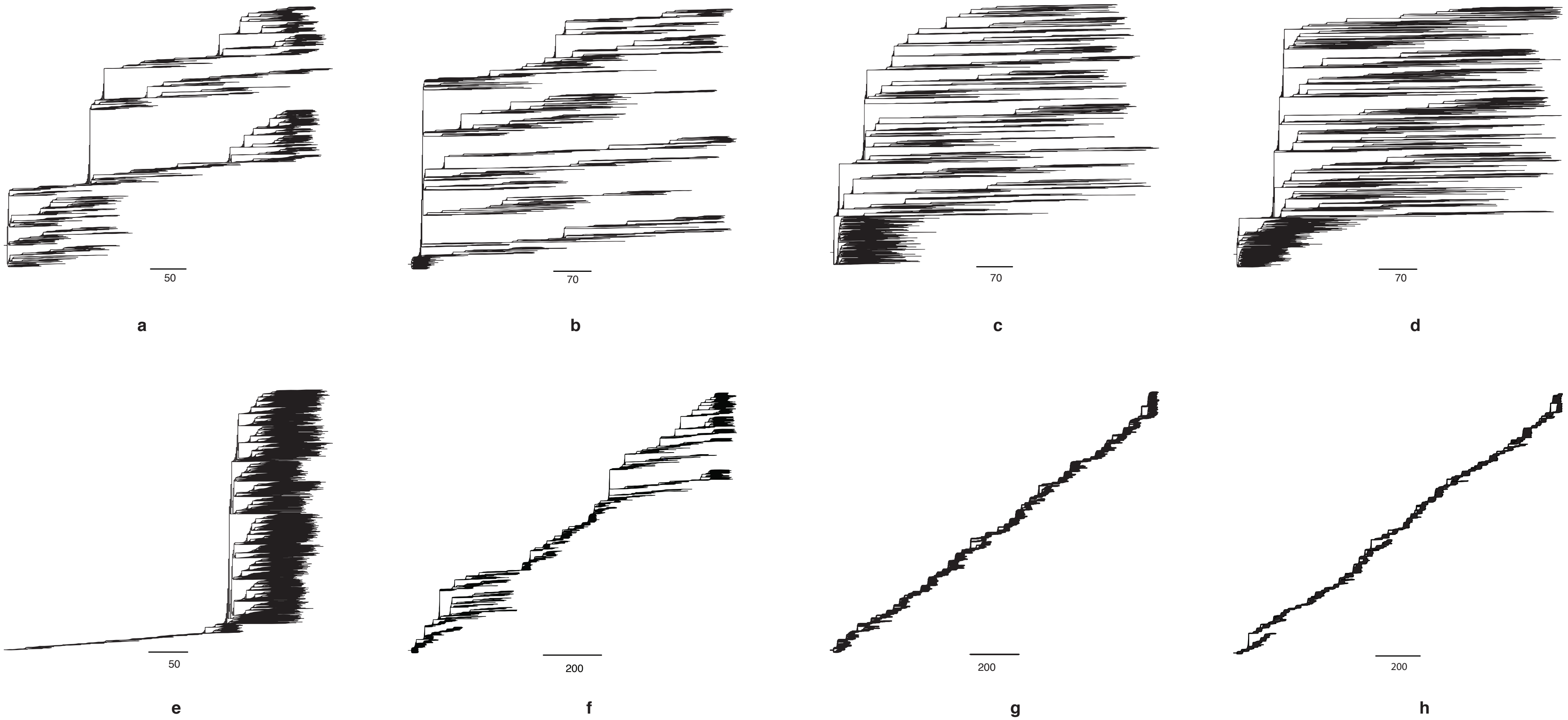

### Supplementary Figure S26

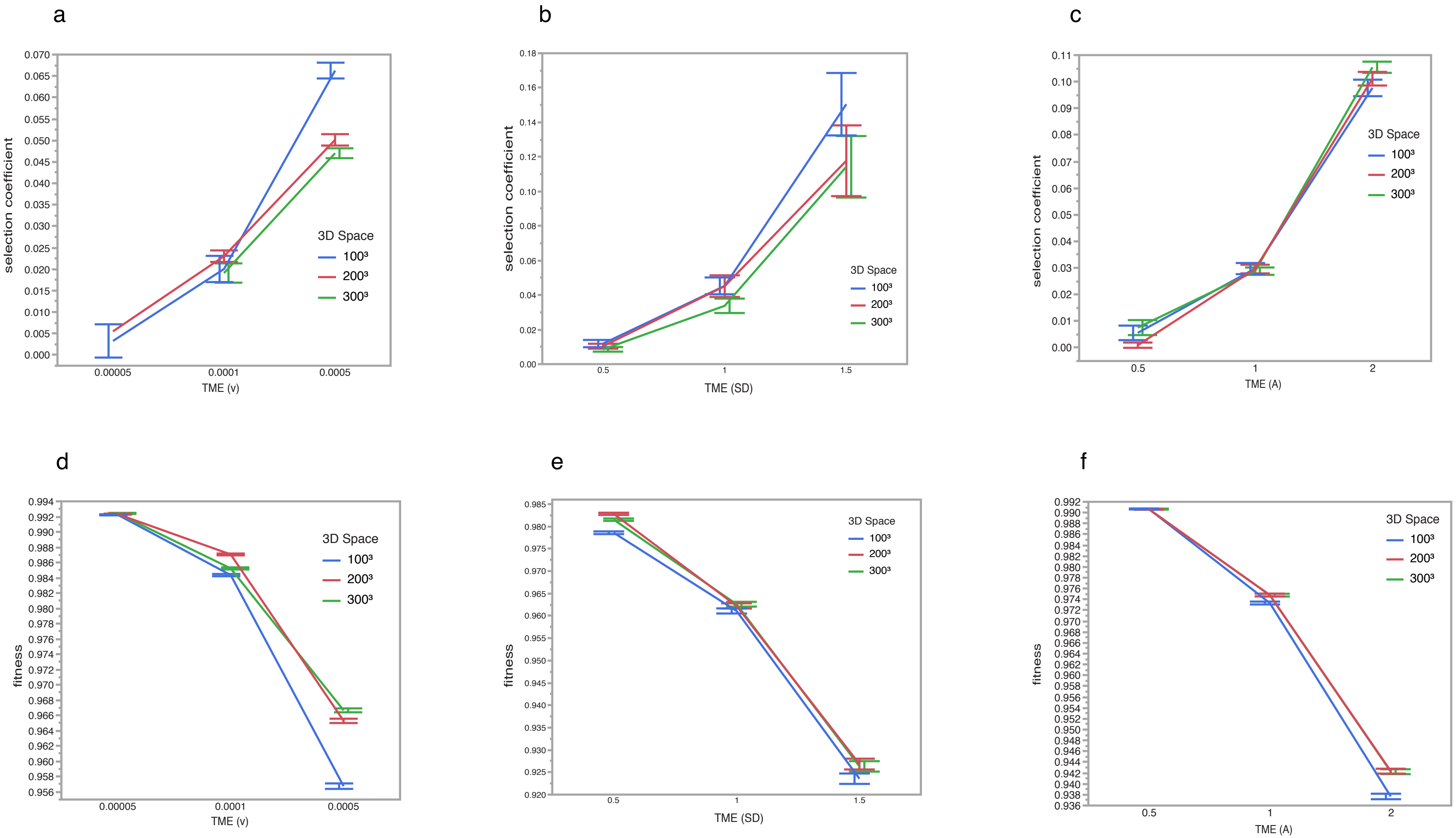

### Supplementary Figure S27

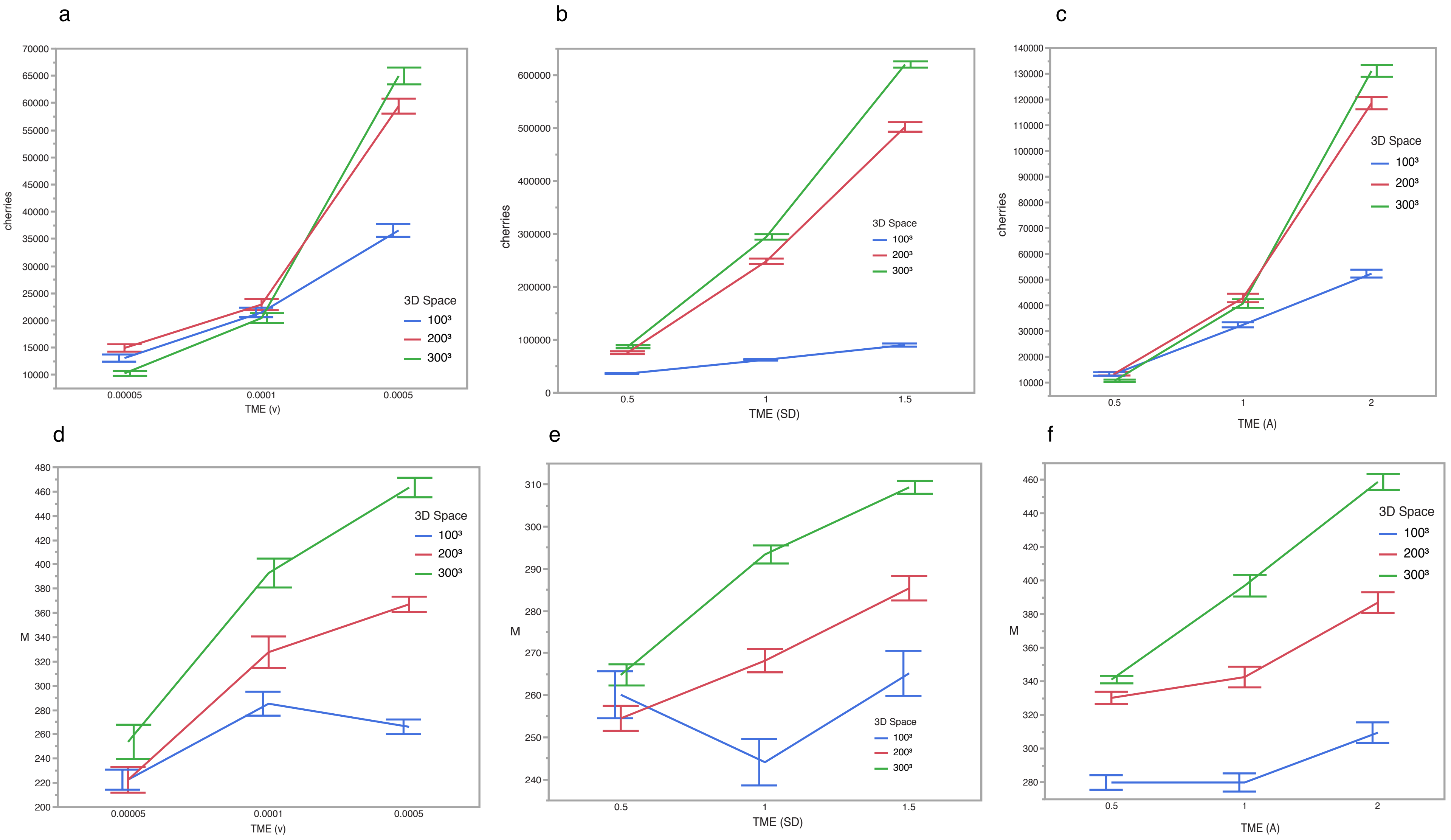

### Supplementary Figure S28

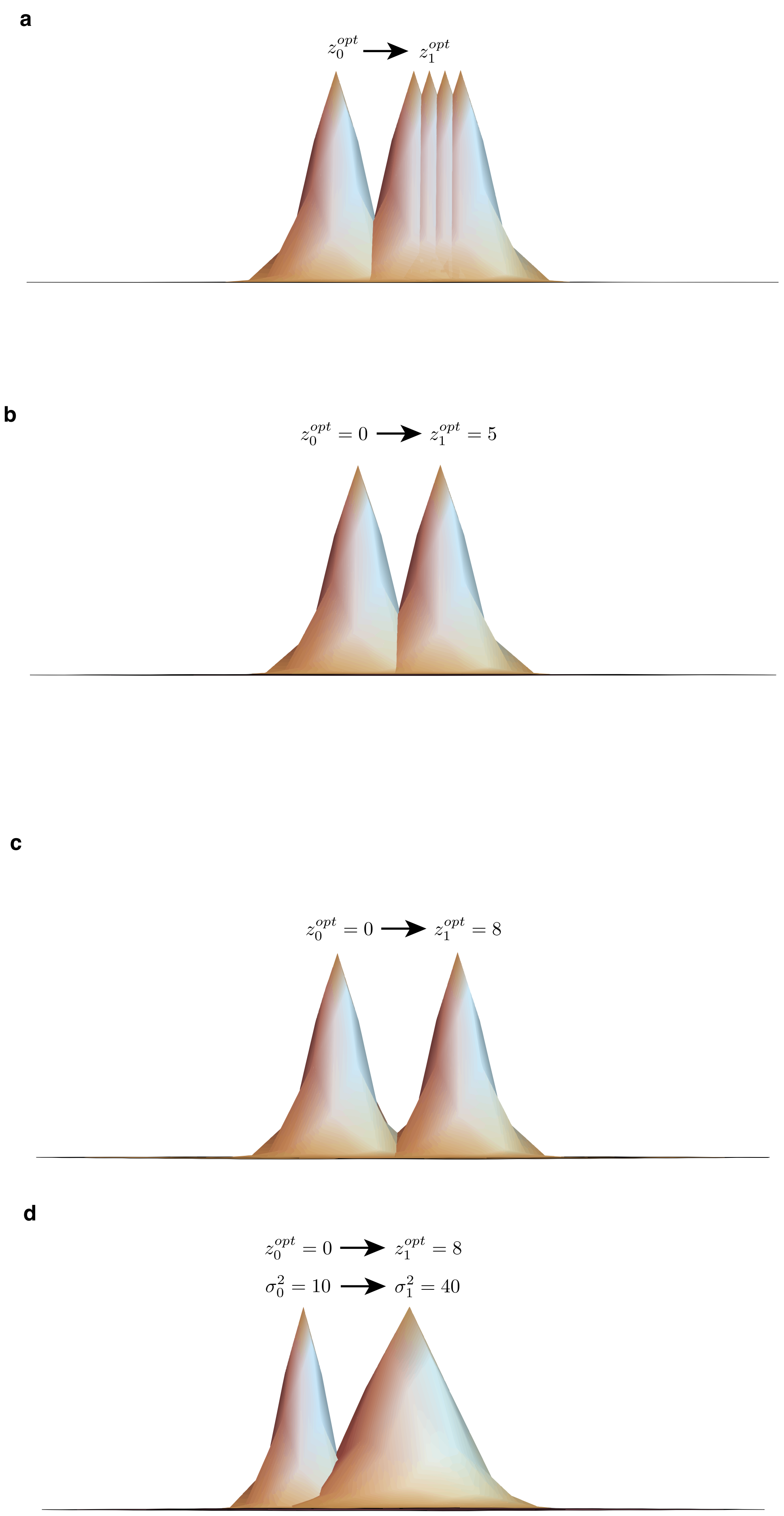

### Supplementary Figure S29

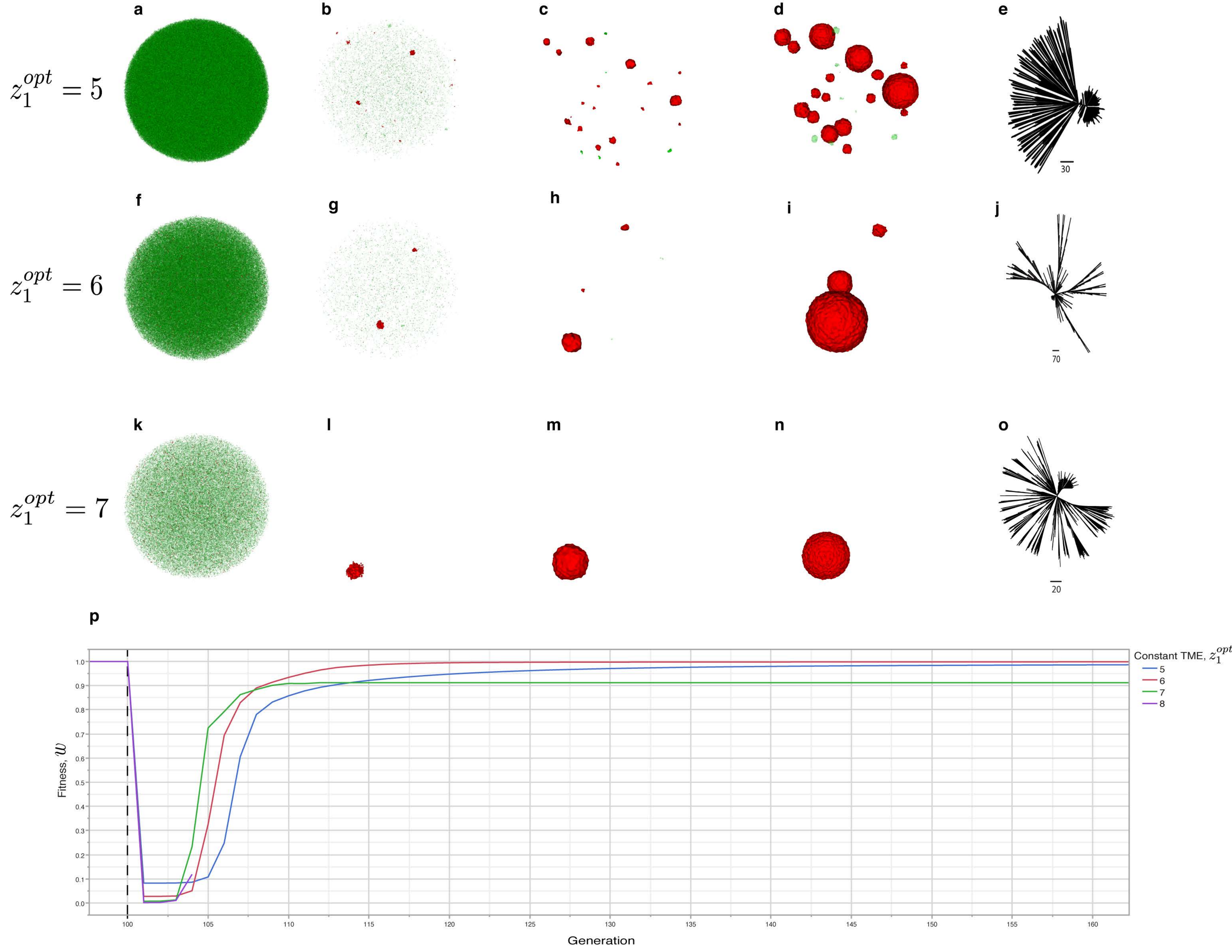

### Supplementary Figure S30

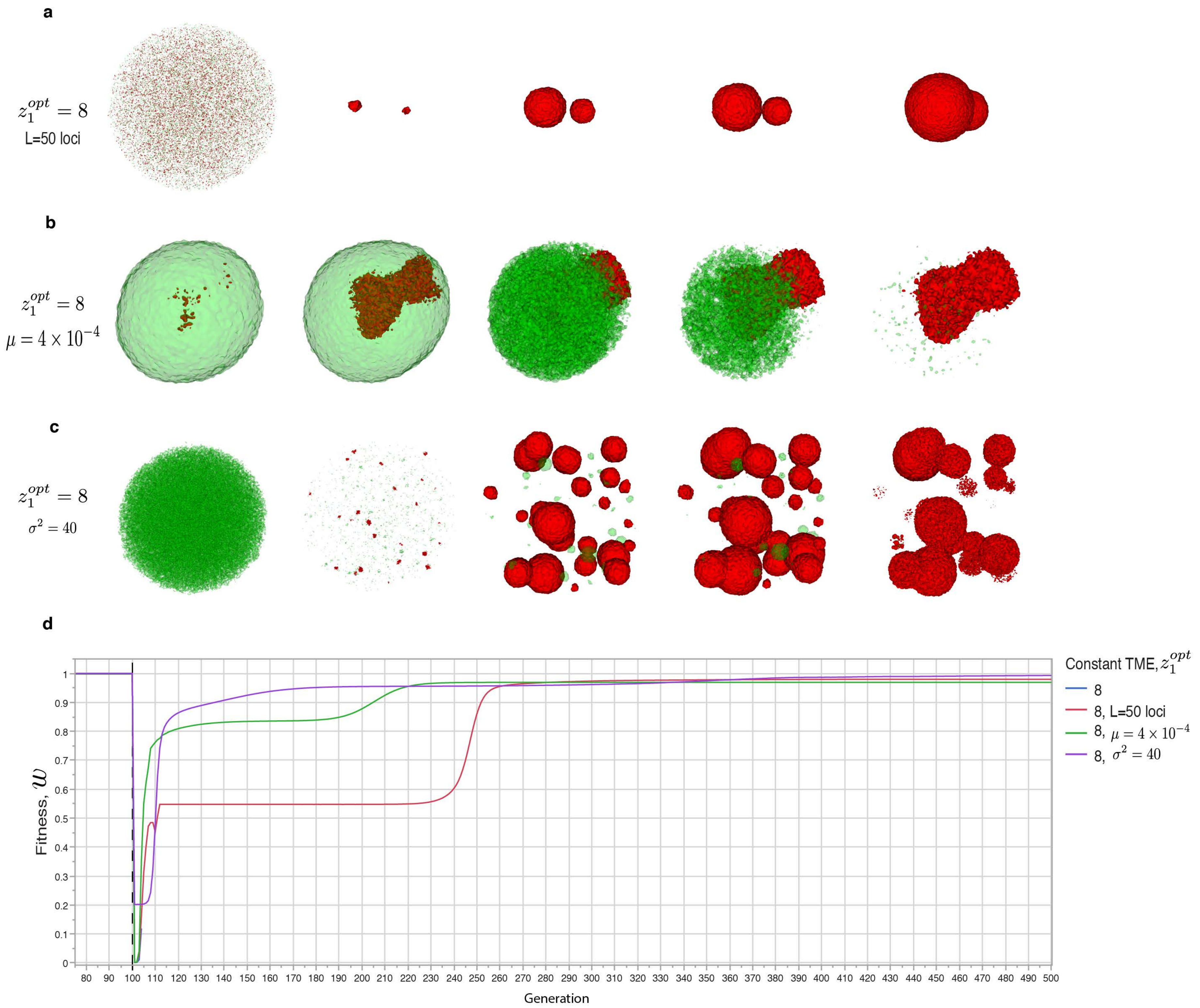

### Supplementary Figure S31

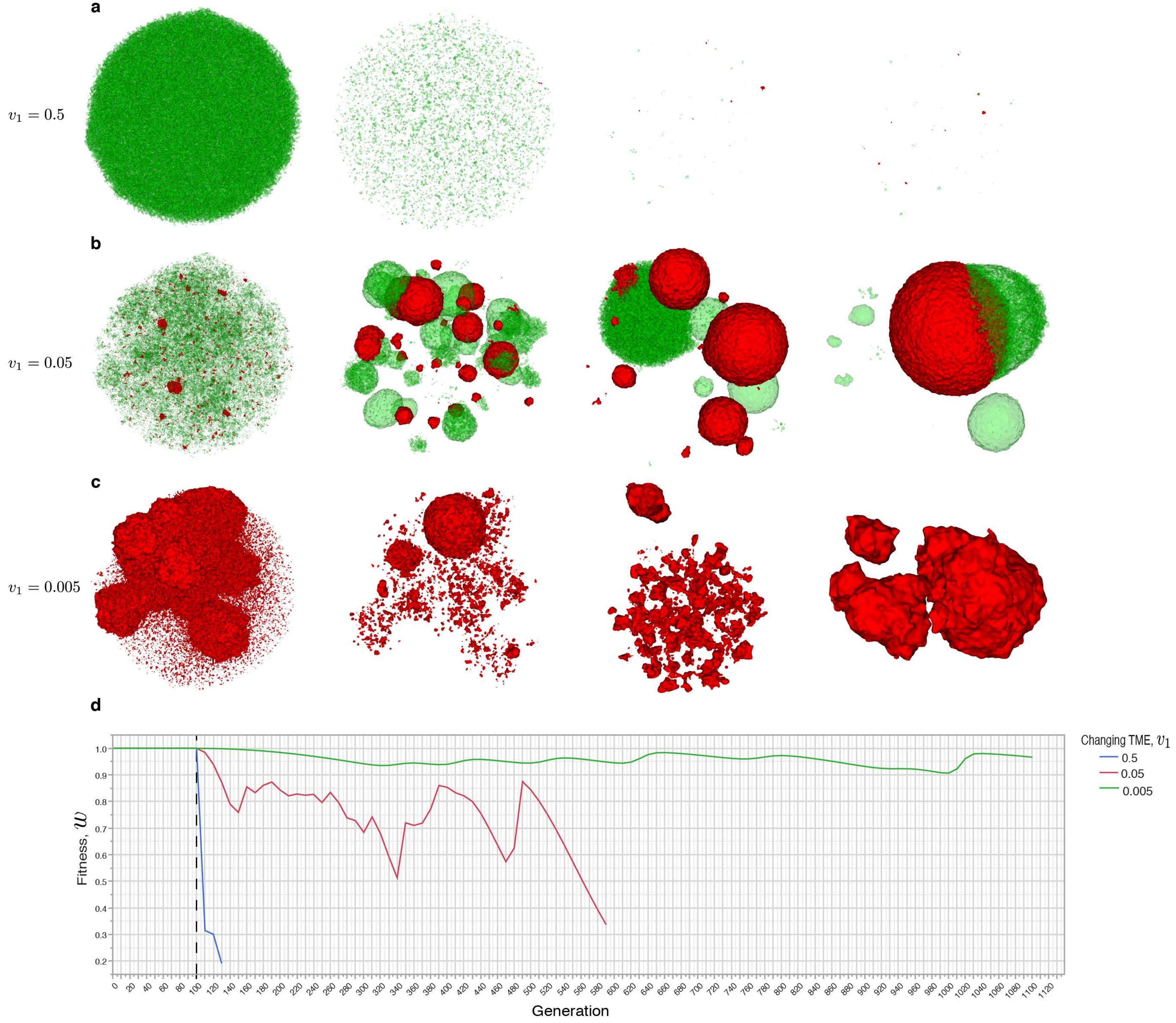
